# Functional Calmodulin States are Selected from an Electrostatically Tuned Free Energy Landscape

**DOI:** 10.64898/2026.01.18.700137

**Authors:** Busra Tayhan, Sila Horozoglu, Ali Rana Atilgan, Canan Atilgan

**Author notes:** **Correspondence:** Canan Atilgan, Faculty of Engineering and Natural Sciences, Sabanci University, Tuzla 34956 Istanbul, Turkey.

## Abstract

Calmodulin (CaM) is a versatile calcium-binding protein whose structural flexibility enables regulation of diverse cellular processes. Capturing its full conformational landscape remains challenging due to high energy barriers between states. Here we employ well-tempered metadynamics simulations using key collective variables to explore CaM conformations under calcium-bound and calcium-free states at physiological and low salt concentrations. We identify four principal conformations that shift in population depending on calcium binding and ionic strength. Calcium binding favors compact states, while low salt conditions flatten the energy landscape, facilitating transitions, but also causing kinetic trapping due to salt-bridge interactions. Comparison with experimental CaM–protein complexes reveals that target binding stabilizes extended conformations distinct from minima accessible to free CaM. These findings elucidate how calcium and ionic environment orchestrate CaM’s conformational dynamics, enhancing understanding of its functional adaptability in cellular calcium signaling.

## Introduction

Calcium (Ca^2+^), a ubiquitous secondary messenger,^1^ plays a crucial role in regulating cellular processes such as muscle contraction, exocytosis, gene expression, apoptosis, and metabolism. Its homeostasis is maintained by the coordinated exchange between cellular compartments^2^ which balance storage, signaling, and energy production. Organelles such as the endoplasmic reticulum,^3,4^ Golgi apparatus,^5^ mitochondria,^4,6^ and the plasma membrane^1^ coordinate its movement to maintain homeostasis and support essential cellular functions. Despite its necessity for cellular function, excessive calcium accumulation can be toxic.^7^ To prevent toxicity and maintain Ca^2+^ signaling, the concentration of this divalent ion is tightly regulated by calcium-binding proteins.

Although many proteins interact with Ca^2+^, only a few show high-affinity and specific binding.^8^ Among them, calmodulin (CaM) stands out for its conformational flexibility and functional versatility.^9^ CaM is a small, acidic, and highly conserved Ca^2+^-binding protein present in all eukaryotes.^10^ It has 148 residues and contains two globular domains connected by a linker (**Figure 1A**). The N-terminal and C-terminal domains span residues 1–68 and 92–148, respectively, each capable of binding two Ca^2+^ ions through EF-hand motifs.^11,12^ The negatively charged residues within these motifs create strong affinity for Ca^2+^ whose binding stabilizes the protein while triggering conformational changes.^13^ The flexible central helix formed by residues 69–91 connecting the two domains imparts controlled flexibility to the protein^11^ which adopts a wide range of conformations.^14^ This flexibility allows CaM to adjust its structure depending on its binding partners.^15^ Ca^2+^ binding reorganizes the domains,^16,17^ either bringing them closer together or moving them apart, directly shaping interactions with target proteins.^13,18^ Although many interactions depend on Ca^2+^, CaM can also bind certain targets in its calcium-free form, reflecting its structural versatility.^19,20^ It can adopt a range of conformations from extended to compact^21^ and engage with binding partners either through both domains or through a single lobe.^22^ This adaptability makes CaM a central regulator of calcium signaling and a mediator of diverse cellular processes.^22^

**Figure 1.**
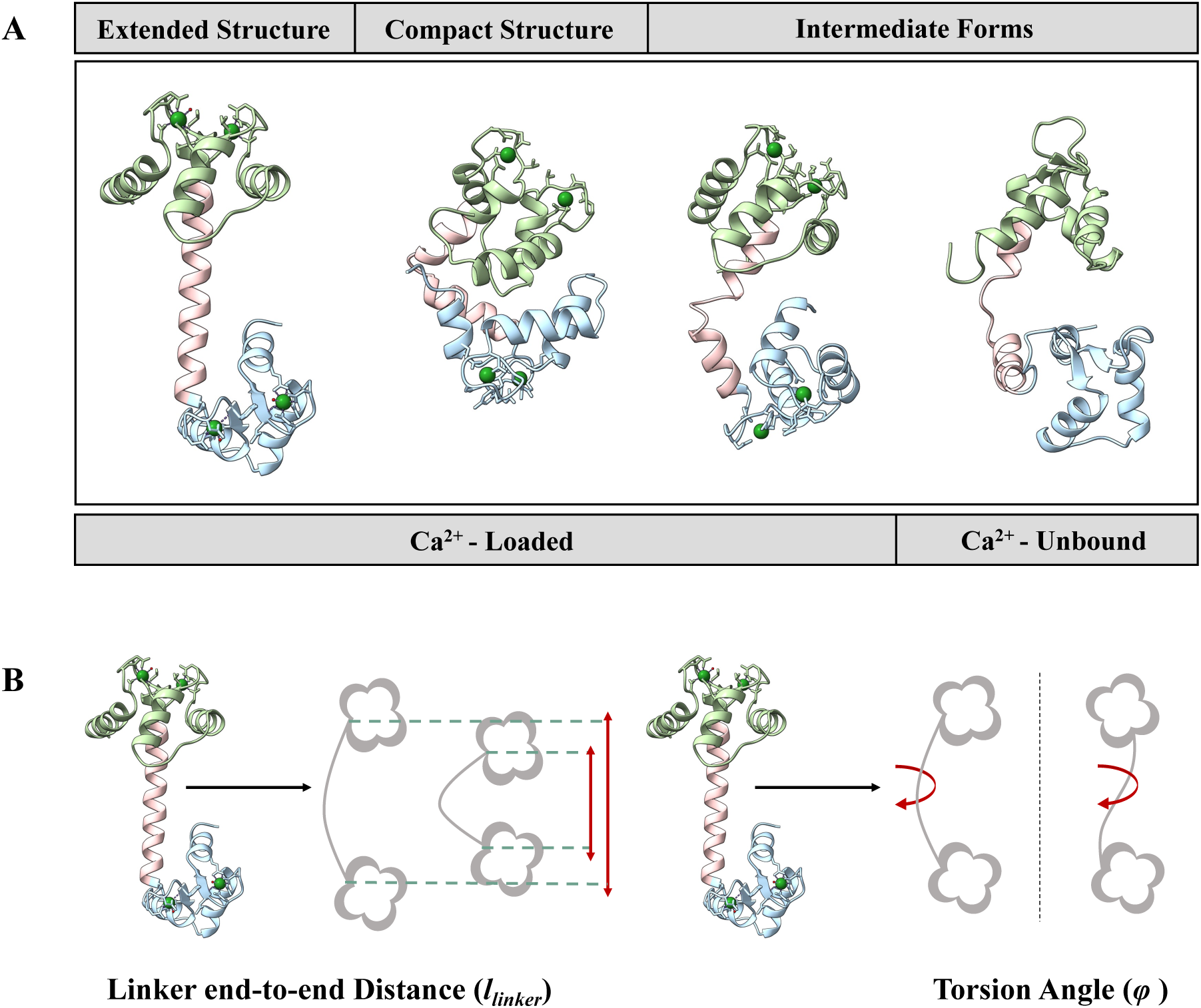
**(A)** Representative three-dimensional structures of CaM. Ca^2+^-loaded conformations include the extended (PDB ID: 3CLN) and compact (PDB ID: 1PRW) X-ray structures. Intermediate forms are captured in the ensembles of NMR structures, e.g. by the Ca^2+^-loaded CaM (PDB ID: 2K0E) and by the calcium-free CaM (PDB ID: 6Y95); two such intermediate conformers from the respective NMR structures are exemplified here. Ca^2+^ ions shown in green; EF-hand motif residue side chains in licorice representation. **(B)** Schematization of the CVs utilized in this study.

The distinct conformations of CaM have been extensively characterized through NMR and X-ray crystallography in calcium-bound, target-bound, and unbound forms, as well as fluorescence resonance energy transfer (FRET)^23^ and mass spectrometry (MS).^24^ Some representative structures are shown in **Figure 1A**.

In its calcium-loaded, peptide-free state, CaM exhibits both extended and compact conformations. X-ray structures often capture an extended dumbbell-shaped arrangement, where the two domains are connected by an elongated helix exemplified by the 3CLN coded Protein Data Bank (PDB)^25^ structure where the two lobes face opposite sides of the linker in a *trans* arrangement.^11^ On the other hand, alternative crystal structures reveal a more compact form with the lobes positioned closer together, e.g. captured in the 1PRW coded PDB structure where the lobes face the same side of the linker in a *cis* arrangement.^26^ In the extended state, CaM exposes hydrophobic pockets that mediate interactions with calcium-regulated enzymes such as kinases and phosphatases, as well as ion channels including the ryanodine receptor and voltage-gated calcium channels, which are central to calcium homeostasis and excitability.^13,27,28^ Ca^2+^-bound, ligand free NMR structures further emphasize CaM’s intrinsic flexibility, displaying a continuum from extended to compact states (e.g. the 160 NMR ensemble structures in PDB ID: 2K0E).^10^ Calcium binding alone does not confine CaM to a single structure but primes it for dynamic transitions.^10^ When bound to target peptides, such as myosin light chain kinase (MLCK), CaM adopts a compact conformation, wrapping tightly around hydrophobic motifs to mediate specific regulatory interactions.^16,17,28^ In the absence of calcium, *apo* CaM typically adopts an intermediate conformation with partially collapsed lobes that retain flexibility;^15^ these are exemplified in the 30 NMR structures of PBD ID: 6Y95.^29^ This state is expected to prime CaM for recognition of conserved IQ motifs in voltage-gated sodium channels and unconventional myosins,^20^ providing calcium-independent regulation of neuronal signaling and intracellular trafficking.^28^

Experimental evidence from FRET and MS studies further supports the conformational flexibility of CaM. Both techniques indicate that calcium-loaded CaM predominantly adopts a compact conformation, challenging the traditionally assumed dumbbell-shaped structure.^24,30^ MS data also show that this globular shape of CaM allows for stable peptide binding, with interactions occurring across both lobes.^24^ Additionally, FRET experiments reveal that *apo* CaM is more extended compared to the globular Ca^2+^-loaded state and supports the view that it exhibits an intermediate conformation.^30^

Despite extensive experimental and computational studies, fully capturing the conformational landscape of CaM remains a challenge. Experimental techniques such as NMR and X-ray crystallography provide representations of CaM’s different states but fail to reveal the numerous transitions between them. Molecular dynamics (MD) simulations offer dynamic insights into the effect of environment on protein conformation and dynamics for this and other metal binding proteins.^31–33^ However, these are often restricted by limited time scales of the simulations, which are insufficient to sample the rare transitions between distinct conformational states.^34,35^ Given the importance of conformational multiplicity in CaM’s ability to regulate cellular processes, applying enhanced sampling techniques^36–39^ might play a crucial role in bridging this gap.

In this study we employed well-tempered metadynamics (MetaD),^40^ a powerful enhanced sampling method that allows us to efficiently explore the free energy landscape of CaM. MetaD introduces a bias potential along selected collective variables (CVs), allowing the system to escape local minima via iterative deposition of Gaussian hills. Once the CVs are sampled with approximately uniform probability, the simulation is halted and the negative of the accumulated bias reveals the underlying free-energy surface. Well-tempered MetaD refines this approach by gradually reducing bias deposition, preventing overfilling and yielding a more accurate reconstruction of the thermodynamic landscape.^40^ Since convergence and a realistic projection of the true free energy surface strongly depends on the choice of the CVs,^41^ their poor selection can cause inadequate sampling and obscure key conformational changes.^42^

Different types of CVs may be selected depending on how effectively they represent the motions of the system. They may be selected from physics-based reaction coordinates, such as inter-atomic distances, angles, dihedrals, or the radius of gyration of a subset of atoms in the system. Alternatively, CVs can be deduced from MD trajectories, such as principal components (PCs) or time-lagged independent components (TICs).^43^ The advantage of selecting physics-based reaction coordinates is that, when comparing the same system under slightly different conditions, the changes may always be followed intuitively and comparatively.^44^ Conversely, when using PCs or TICs, there is no guarantee that the projections reflect similar phenomena which renders interpreting shifts in the energy landscape tricky.^44,45^ Therefore, we used two physics-based reaction coordinates that were shown to capture the motions of CaM: the rotational motion of its lobes relative to each other and the change in their separation (**Figure 1B**).^31^ These DoFs were defined to project MD trajectories into conformational subspaces to provide a detailed understanding of population shifts among CaM’s conformational states and have also been applied in other contexts.^46^

We leverage well-tempered MetaD in combination with classical MD simulations to map how calcium loading, ionic strength, and the ensuing electrostatic environment shape the conformational free-energy landscape of CaM. Mapping conformations on the landscape captured by our pre-defined CVs allows us to capture rare transitions, identify previously uncharacterized minima, and distinguish thermodynamically favored states from kinetically persistent ones. By integrating these computational landscapes with experimentally determined CaM–target complexes, we show that CaM’s functional conformations are selected from a tunable electrostatic landscape rather than predetermined by its intrinsic minima. This framework provides a mechanistic basis for how CaM adapts its structure to cellular context and target engagement, offering broader insights into how electrostatics and environmental cues regulate protein conformational ensembles.

## Results and Discussion

To dissect how calcium loading and ionic strength shape the conformational landscape of calmodulin, we first examined the extent to which unbiased MD can sample the relevant structural states under each condition with duplicate 1 µs runs (see RMSD profiles in **Figure S1**). As expected, cMD simulations were not efficient in sampling possible *apo*/*holo* conformations of CaM under the different environmental conditions. In all but one case, the linker maintained its original rigid α-helical structure (*l*_linker_ in the range of 30-35 Å), while in all cases the two lobes displayed a repositioning with respect to the initial 3CLN structure traced by the torsional CV, *φ*, from an initial *trans* positioning of 100° to a predominant sampling in the range (−180,0°) (**Figure S2)**. The highest mobility is observed in the H^L^ system, and from the RMSD plots of the individual cMD runs, we can see that one of the two trajectories is responsible for the jump to the compact conformer at ca. 400 ns timepoint (**Figure S1**). Focusing on the RMSD of individual domains, we find that the N-lobe is always more mobile than the C-lobe irrespective of the environmental conditions, more so in the *apo* systems than the *holo* systems. These observations are in agreement with experiments where the fast internal mobility of the N-terminal domain have long been established via NMR experiments,^51^ and its lower calcium affinity has been measured under low salt and physiological conditions.^52^ While cMD simulations concur with the relative flexibilities of the two domains, it is also evident from **Figure S2** that the observed conformers are a small subset of those that are physically available to CaM, represented by the larger dots. To sample the full conformational space available to CaM under these differing conditions, we resort to MetaD simulations in what follows.

### Effects of calcium loading to CaM conformers at physiological salt

The time evolution of the selected CVs during the MetaD runs are displayed in **Figure S3**; we note that the corresponding PMFs (**Figure S4**) are converged by 400 ns, and an additional 200 ns is run to ensure the convergence of the potentials of mean force (PMFs). Interestingly, we find the relative positioning of the two lobes, represented by the torsional CV, *φ*, is sampled on a faster time scale than *l*_linker_. This difference in the time scales is a testament to the difficulty of positioning the two highly negatively charged lobes in proximity and the requirement of finding an facilitating arrangement in the system to reach these compact conformers.

To systematically analyze the free energy landscape, we normalize the results for each system taking the open dumbbell-shaped crystal structure of CaM (3CLN) as a reference. The CVs of the reference structure are calculated and mapped to the PMFs for each system. Energies corresponding to (*l*_linker_, *φ*) values of the reference structure are extracted and set as the zero of the PMF in each case, serving as baselines for normalization. The PMF plots are then shifted to compare all surfaces with respect to this common reference energy level (**Figure S4**); the two dimensional projections of these surfaces are displayed in **Figure 2** where the color bar reflects this standardized range, from −9 to 6 kcal/mol. For reference, positions of the experimental structures are also overlaid onto the surfaces of **Figure 2** where black dots represent NMR structures determined for *holo* (2K0E) or *apo* CaM (6Y95); and teal dots mark crystal structures (3CLN and 1PRW).

**Figure 2.**
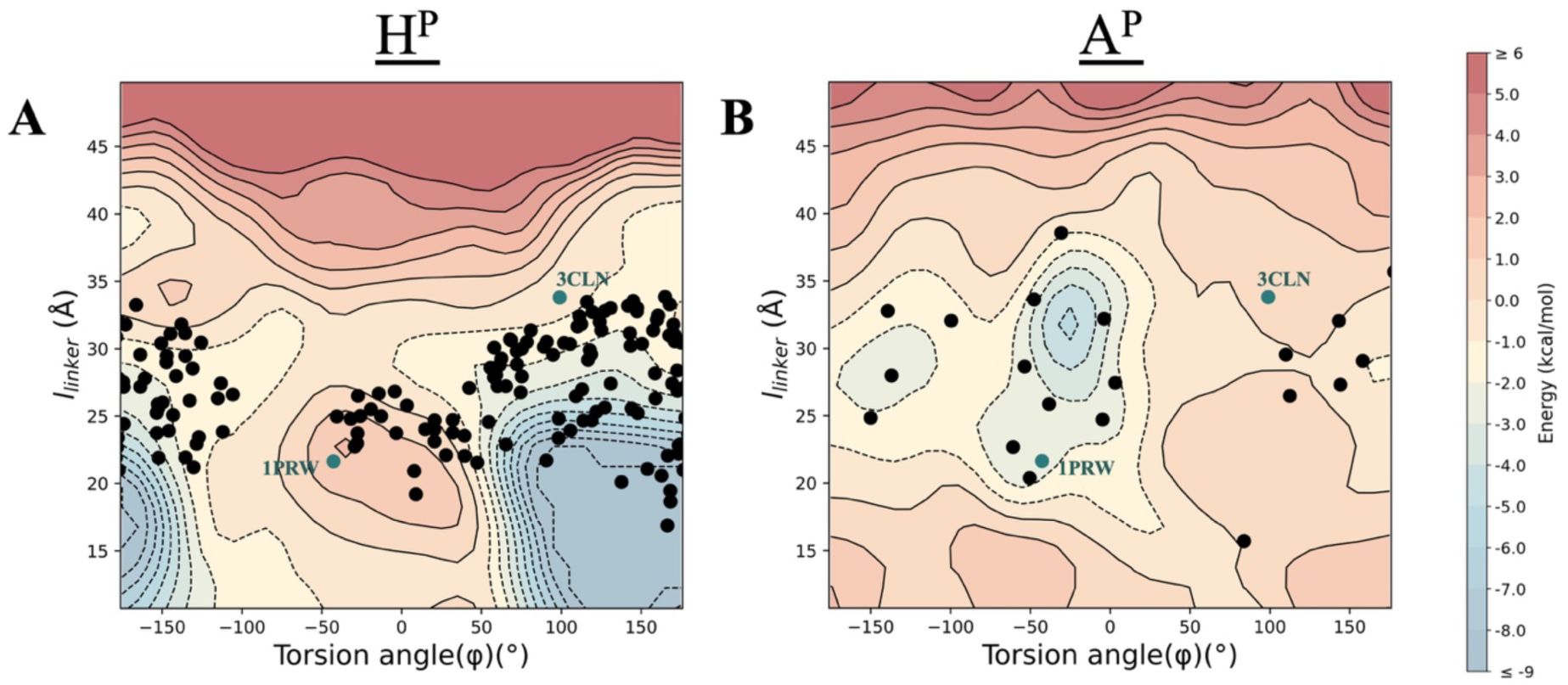
Free energy surface of CaM obtained from MetaD in physiological salt concentrations; **(A)** *holo* CaM and **(B)** *apo* CaM. Black dots represent NMR (2K0E for *holo*, 6Y95 for *apo* CaM) structures while teal dots are labeled with the PDB code of the crystal structures they represent.

The PMF of H^P^, obtained from MetaD (**Figure 2A**) reveals a single prominent minimum characterized by the positioning of the two lobes mostly facing each other, quantified by the wide distribution around *φ*=150° and a linker distance of up to 20 Å. We label this conformation where the N- and C-lobes face opposite sides *trans*-compact whose minimum is located −15.3 kcal/mol with respect to the 3CLN structure. Compact states are also observed in NMR structures of *holo* CaM (2K0E); some of which even more compact than the 1PRW X-ray structure. Mass spectrometry experiments show that CaM predominantly adopts compact conformations in fully Ca^2+^ loaded CaM,^24^ supporting our MetaD findings. While few of the 160 representative NMR-deduced structures mapped onto the PMF are compact, the majority occupy the regions that are defined by the basin mapped by MetaD. We note that these experimental data were collected^53^ at 100 and 10 mM and collated to generate the 2K0E conformers.^10^ We will discuss the PMF obtained under low salt conditions in the next subsection.

In the A^P^ system (**Figure 2B**), a major conformer with the two lobes facing the same side of the linker (centered around *φ* = −30°) and a minor conformer where they face the opposite side (centered around *φ* = −150°) are found. In both minima, the linker maintains its straight helical initial conformation with *l*_linker_ > 25 Å. We term these respective conformers *cis*-extended and *trans*-extended, where the minima are at −7.1 kcal/mol and −3.1 kcal/mol with respect to the reference position. The structures from the NMR dataset of *apo* CaM (PDB ID: 6Y95), all collected at 105 mM, align closely with our PMF basins. Thus, we conjecture that the *apo* structure under physiological conditions displays a rigid yet rotatable linker whose axial rotation allows the N- and C-terminal calcium-binding lobes to adopt either facing (major conformer) or opposing (minor conformer) orientations. Such reorientation effectively exposes both EF-hand domains to the surrounding solvent, thereby facilitating efficient sampling of the local Ca^2+^ environment. This intrinsic positional freedom is consistent with CaM’s role as a high-sensitivity Ca^2+^ sensor whereby it permits both lobes to act semi-independently so that the protein can rapidly engage available ions from multiple spatial directions.

### Effects of low-salt conditions on CaM conformations

While CaM is not typically found in low-salt conditions, but rather in intracellular environments of ~150 mM salt, there is ample literature where its structure and biochemistry is scrutinized under the former conditions. This is mainly because Ca^2+^ binding and domain–domain contacts in CaM depend heavily on electrostatics. Reducing ionic strength with low-salt buffers minimizes charge screening, amplifying electrostatic effects. This helps reveal details about Ca^2+^ binding cooperativity, interdomain communication, and long-range conformational coupling that are otherwise masked at physiological salt. Moreover, at low ionic strength, fewer ions compete with Ca^2+^ for acidic residues, making the intrinsic binding constants easier to measure. Classic studies used these conditions to separate N- and C-lobe affinities unambiguously^52^ and led to the conclusive result that Ca^2+^ binding is cooperative within each of the domains while there is no indication of cooperativity between the domains.

Notably, local ionic microenvironments, e.g., near negatively charged membranes or within protein complexes, can transiently reduce the effective electrostatic screening experienced by CaM, even when the bulk ionic strength remains physiological. In such contexts, counterion accumulation, co-ion exclusion, and geometrical confinement enhance long-range electrostatic interactions in ways that resemble low-salt conditions *in vitro*. Thus, low-salt experiments can serve as controlled physical probes for uncovering coupling mechanisms and long-range interactions that remain operative, though partially masked, under normal intracellular ionic strengths. To investigate the effect of decreasing monovalent ions in the environment on the conformations of CaM, we performed a 600 ns-long MetaD simulations of *apo* and *holo* CaM mimicking low salt conditions (only K^+^ present in the periodic cell) (see **Table 1** under **Methods**). We then applied the same normalization and scaling procedure as before to systematically analyze their free energy landscape with respect to the position of the canonical 3CLN conformation which is placed at 0 kcal/mol as reference.

**Table 1.** Summary of systems simulated.

| System label* | Equilibrated box size (Å<sup>3</sup>) | Salt ions (K<sup>+</sup>/Cl<sup>-</sup>) | Ionic strength (mM) | Simulation time (cMD/MetaD/cMD) |
| --- | --- | --- | --- | --- |
| H<sup>P</sup> | 65.3×82.4×61.8 | 43 / 28 | 177 | 2×1μs / 600 ns / 4×0.2 μs |
| A<sup>P</sup> | 65.1×82.3×61.6 | 51 / 28 | 199 | 2×1μs / 600 ns / 4×0.2 μs |
| H<sup>L</sup> | 65.3×82.4×61.7 | 15 / 0 | 38 | 2×1μs / 600 ns / 4×0.2 μs |
| A<sup>L</sup> | 65.3×82.5×61.8 | 23 / 0 | 57 | 2×1μs / 600 ns / 4×0.2 μs |

\* H/A labels indicate *holo/apo* CaM; superscripts P/L indicate physiological/low salt conditions.

Compared to their physiological salt counterparts, the PMF of both H^L^ and A^L^ exhibits shallower minima separated by lower energy barriers (**Figure 3**). H^L^ maps a large basin on the conformational surface; but we can distinguish two lower energy regions, each corresponding to a specific structural conformation (**Figure 3A**). The first is in the wide basin spanning torsion angle in the range [-40°, 60°] and a linker distance range of 16–26 Å; while this region has three closely spaced local minima, together we label it the *cis*-compact conformation located at −4.1 kcal/mol global minimum. The second distinct region has a torsion angle centered on 180° and with a linker distance in a limited range of 18–21 Å, labelled *trans*-compact located at −3.0 kcal/mol. We also see that the regions of extended *cis* and *trans* conformations are accessible in this ionic strength as relatively low-lying energy regions. Most of the representative NMR conformers deposited under PDB ID 2K0E populate the same regions of the free-energy surface identified in our low-salt simulations. As noted earlier, these experimental structures were derived from datasets acquired at two distinct ionic strengths (10 and 100 mM), preventing unambiguous assignment of individual conformers to a specific condition. Nonetheless, their ensemble distribution is consistent with the conformational basins resolved by our MetaD analysis, indicating that when we consider our H^P^ and H^L^ simulations together, they faithfully capture experimentally accessible conformations.

**Figure 3.**
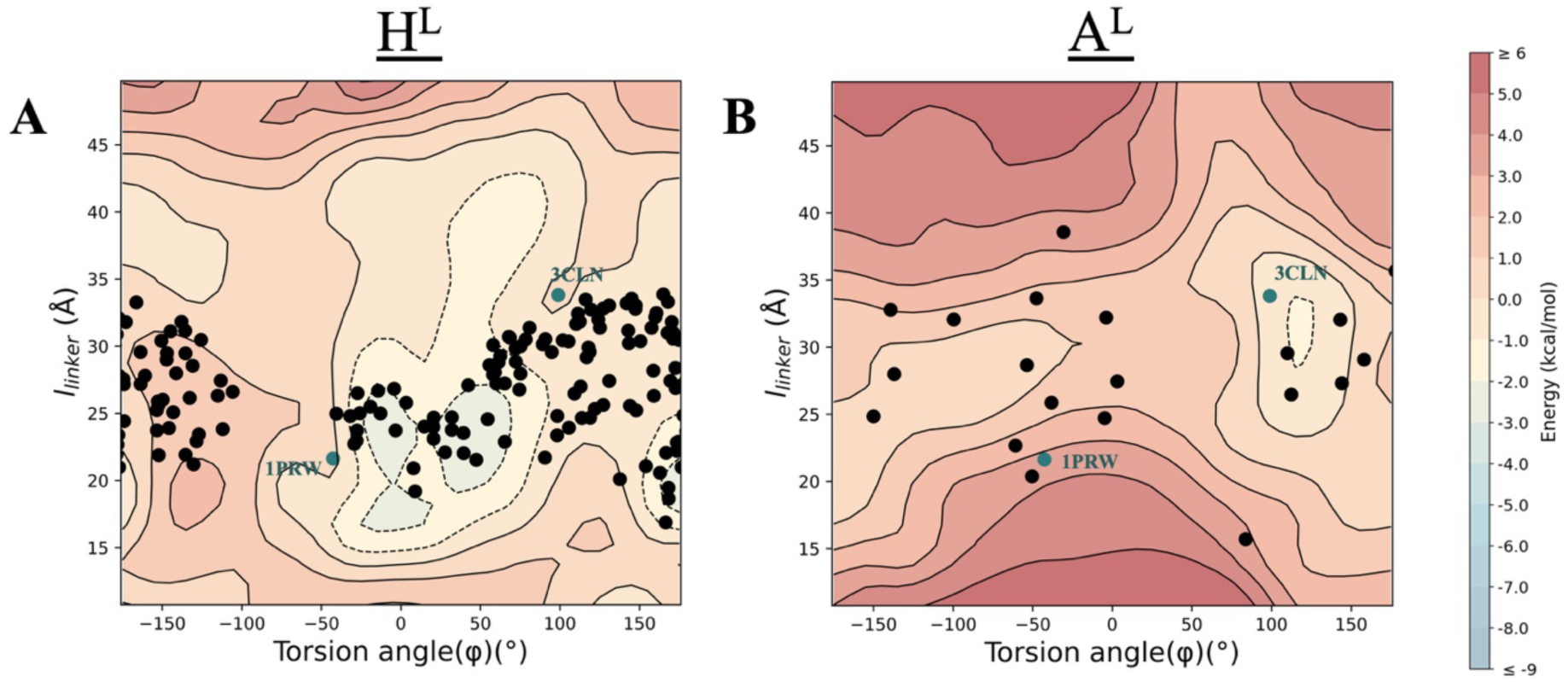
Free energy surface of CaM obtained from MetaD in low salt; **(A)** *holo* CaM and **(B)** *apo* CaM. Dots represent the same NMR and crystal structures as in Figure 2.

The A^L^ PMF contains a single minimum characterized by dihedral angles > 100° and linker distances centered around 32 Å, corresponding to a *trans*-extended conformation located at −1.6 kcal/mol (**Figure 3B**). This minimum is markedly shallower than those in the other systems, reflecting a softer energetic-landscape in which *apo* CaM adopts an open, dumbbell-like architecture reminiscent of the canonical *holo* 3CLN conformation (**Figure 1**). It is worth noting that our *apo* simulations span distinct ionic strengths of 199 mM for A^P^ and 57 mM for A^L^, while the experimental *apo* NMR ensemble (PDB 6Y95) was acquired at an intermediate ionic strength of 105 mM. Although this complicates direct comparison, the experimental structures fall within the broader conformational regions predicted by our A^P^ and A^L^ surfaces, supporting the robustness of the simulated landscapes.

These results show that lowering the ionic strength substantially softens the conformational energy landscape of CaM, promoting facile interconversion among compact and extended states in both the *holo* and *apo* forms. For *holo* CaM, reduced ionic screening diminishes the dominance of the *cis*-compact basin and stabilizes additional compact substates that are separated by only small energetic differences, facilitating rapid interdomain rearrangements. For *apo* CaM, the shallow *trans*-extended minimum reflects an intrinsically flexible, weakly biased landscape that readily accommodates repositioning of the two lobes around an elongated central linker. Such energetic flattening is consistent with a scenario in which reduced ionic strength enhances long-range electrostatic coupling while lowering the penalties associated with domain reorientation. Although CaM typically operates at physiological salt concentrations, local electrostatic microdomains, such as those near membrane surfaces, within crowded assemblies, or in regions of fluctuating ion flux, may transiently reduce effective screening, creating environments that electrostatically approximate low-salt conditions. Our findings therefore suggest that CaM retains the capacity to explore a broadened conformational repertoire within such environments, providing a biophysical basis for its rapid structural adaptability during target engagement and signaling.

### Stability and Kinetic Accessibility of Minima Assessed by Classical MD Simulations

To determine whether the conformational minima identified through MetaD simulations correspond to kinetically stable states or merely reflect transiently sampled basins, we next examine their dynamical behavior using unbiased cMD simulations. While MetaD provides a comprehensive reconstruction of the free-energy landscape, the addition of bias accelerates barrier crossing and may allow sampling of states that are not dynamically stable under equilibrium conditions. cMD therefore offers a complementary view by testing whether these minima persist, interconvert, or dissipate when the system evolves solely under the physical force field. To this end, we extract representative structures from the principal MetaD basins across all systems, subject them to minimization and equilibration, and monitor their spontaneous dynamics projected onto our selected CVs. This approach allows us to evaluate the robustness of the MetaD-derived minima, identify potential kinetic traps, and dissect the role of ionic strength and Ca^2+^ loading in stabilizing or destabilizing distinct conformational states.

We identify four representative conformers from the MetaD simulations each representing the structures sampled under the various conditions (**Figure 4A**). Going counterclockwise the quadrants of our coarse-grained conformational space, **I** is a *trans*-extended, **II** is a *cis*-extended, **III** is a *cis*-compact and **IV** is a *trans*-compact conformer. Interestingly, while each of these dominates one of the conditions (**I** ↔ A^L^; **II** ↔ A^P^; **III** ↔ H^L^; **IV** ↔ H^P^), they may still be sampled in more than one condition. In particular, the shallow energy landscape in low salt conditions (**Figure 3**) is expected to lead to easy passage between the minima.

**Figure 4.**
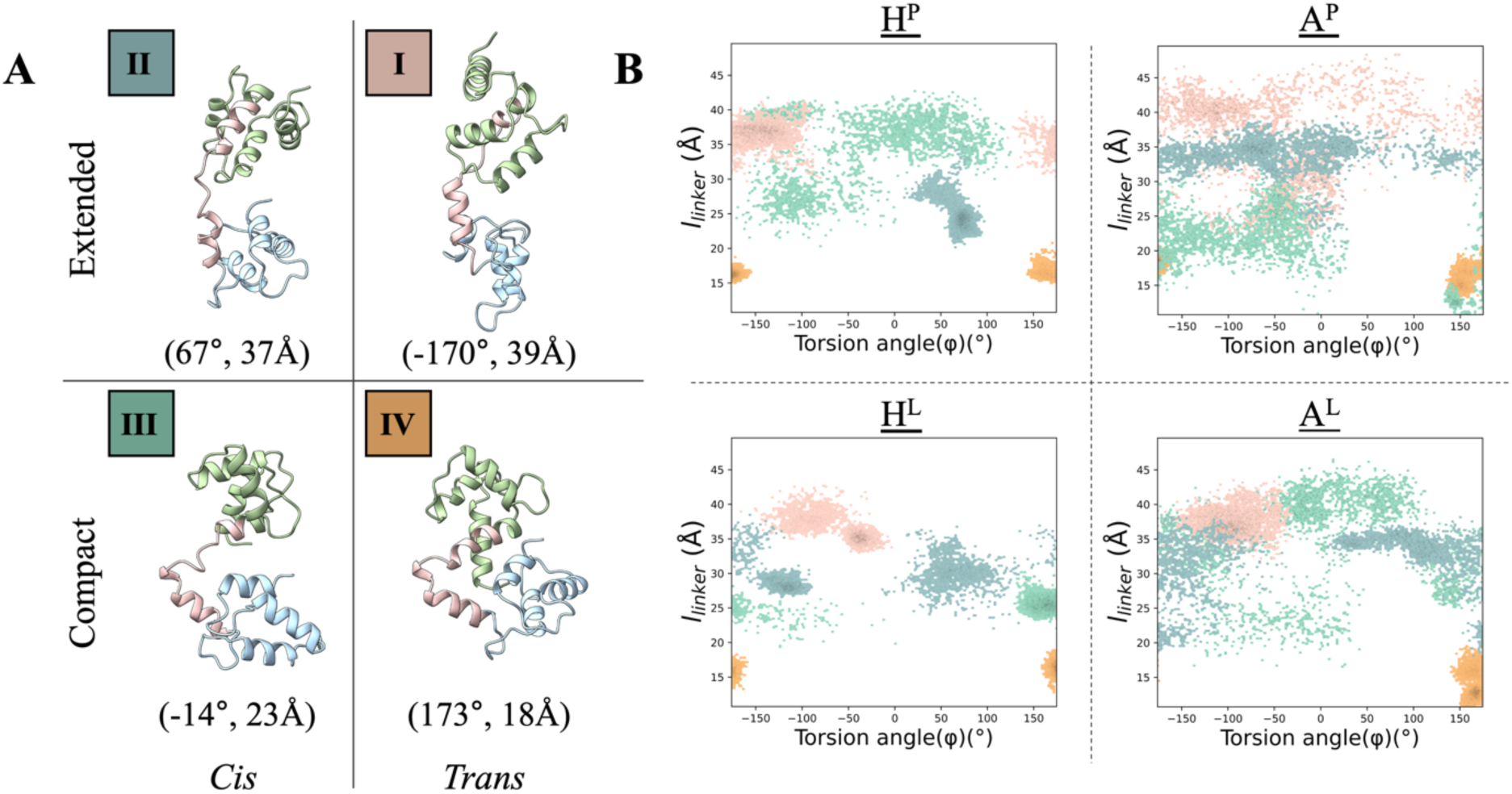
**(A)** The four conformers selected from the PMFs. Their three-dimensional structures and the specific values of the (*φ*, *l*_linker_) are displayed. The color labels are **I**=pink, **II**=teal, **III**=mint, and **IV**=ochre. **(B)** The 4 × 200 ns time points visited under each condition by each conformer are displayed on the CVs with the corresponding colors.

We select snapshots corresponding to these four local minima from our H^P^ MetaD run and then subject them to the appropriate conditions as described under the Methods section to reproduce 200 ns-long cMD simulations for each setup. The coordinates so obtained are projected onto the two-dimensional space in **Figure 4B**, color coded according to the initial structure I-IV.

We make a couple of general observations from these simulations: First, we find that the *apo* systems display a larger conformational freedom than their corresponding *holo* counterparts in the same salt environment. This is to be expected as *apo* CaM lacks the Ca^2+^-mediated intradomain stabilization that normally rigidifies the EF-hand motifs, thereby weakening interhelical packing and permitting larger-amplitude motions of the two lobes relative to the linker. This observation was not fully recovered in the original 2×1 µs cMD simulations which all started from the 3CLN structure (**Figure S2**). Therein, although we observed this extra flexibility in the overall RMSD profiles in the *apo* forms compared to the *holo* forms in **Figure S1**, this flexibility did not usually translate into sampling of multiple conformers. Our second general observation is that, in contrast to the other conformers, **IV** exhibits striking kinetic rigidity; it remains trapped near its *trans*-compact conformation in every condition, including those where it is not a thermodynamic minimum.

Beyond these trends, in the H^P^ system, three of the four conformers (**I**, **II**, and **IV**) remain tightly localized during the cMD trajectories, consistent with their positions in stable or moderately stable regions of the H^P^ free-energy landscape (**Figure 2**). Conformer **IV**, which corresponds to the global *trans*-compact minimum of the H^P^ PMF, is the most confined, but conformers **I** and **II** also remain in relatively well-defined regions. While **II** is drawn to a more compact form within the 200 ns time frame of the simulations, towards the edge of the large *trans*-compact global energy minimum, **I** samples *trans*-extended conformers in our 200 ns window of observations. In contrast, **III** samples a broad, diffuse distribution because the *cis*-compact geometry lies near a local maximum of the H^P^ PMF; lacking stabilizing curvature, it experiences no restoring force, and small thermal fluctuations are sufficient to drive the system away from the initial configuration. This behavior underscores that in H^P^, only the *trans*-compact basin is deeply stabilized, whereas other compact arrangements, particularly the *cis*-compact geometry, are energetically unfavorable and kinetically unstable.

In the A^P^ system, conformers **I-II-III** exhibit broad sampling across the two-dimensional conformational space (**Figure 4B, upper right**), reflecting the inherently flexible nature of *apo* CaM under physiological ionic strength. The MetaD PMF for A^P^ shows a single, moderately deep basin defined by the torsion angles in the range [−50°,+20°] and linker distances of 28–32 Å, corresponding to the *trans*-extended state (**Figure 2**), and this agrees with the dense band of cMD sampling in the same region. These conformers diffuse extensively across torsion space, indicating that the A^P^ landscape contains shallow energetic curvature and permits facile reorientation of the lobes around a relatively pliable linker. Conformer **IV**, although thermodynamically disfavored in A^P^, remains kinetically confined to its compact starting region, producing a narrow sampling cluster near (φ ≈ 150°, *l*_linker_ ≈ 15–20 Å). This behavior reflects strong intra-protein electrostatic clamps, which persist even in the absence of Ca^2+^.

In the H^L^ system, the cMD trajectories (**Figure 4B, lower left**) reflect the shallow, gently partitioned topology of the PMF, in which the deepest region is the broad *cis*-compact triple-minimum basin (−4.1kcal/mol), accompanied by a secondary *trans*-compact minimum located only 1.1 kcal/mol higher (−3.0 kcal/mol) (**Figure 3A**). **IV** remains confined to this latter *trans*-compact well. By contrast, the other three conformers undergo basin-to-basin relaxation: **I**, initialized in the *trans*-extended quadrant, migrates toward the *cis* region with a somewhat shortened *l*_linker_. **II** spreads widely in both torsion angle and linker distance, sampling portions of the broad triple-minima basin. **III**, although initialized in the *cis*-compact region, rapidly drifts towards the *trans*-compact basin. Because the *cis*-compact region in H^L^ does not form a single sharply defined well but instead consists of three adjacent shallow minima separated by barely perceptible ridges, conformers readily slide toward neighboring basins, including the slightly higher-energy *trans*-compact state. These transitions, observed within the limited 200 ns time scale of the simulations, demonstrate that in low salt, the *holo* landscape is dominated by shallow internal structuring, allowing conformers to flow into alternative minima with minimal energetic resistance.

In the A^L^ system (**Figure 4B, lower right**), all starting conformers except **IV** disperse widely across the two-dimensional conformational space, consistent with the extremely shallow, weakly structured topology of its PMF (**Figure 3B**). The PMF displays a single broad minimum centered around φ ≈ 100–140° and linker distances of 28–32 Å, corresponding to a *trans*-extended geometry, accompanied by only very gentle curvature elsewhere. Conformers **I**, **II**, and **III** rapidly lose memory of their initial states and diffuse throughout the extended region of the landscape, sampling a broad continuum of torsion angles and linker distances without settling into a distinct basin. This behavior reflects the absence of Ca^2+^-mediated intradomain stabilization, combined with reduced ionic screening, which together render *apo* CaM highly flexible and prone to large-amplitude domain reorientation. Conformer **IV**, although initiated in the *trans*-compact corner of the landscape, remains kinetically trapped near its starting position, but only because its compact geometry forms a self-stabilizing electrostatic cluster; this trapping does not correspond to any meaningful minimum on the A^L^ PMF. Overall, the A^L^ simulations demonstrate that low ionic strength amplifies the inherent structural pliability of *apo* CaM, yielding a nearly flat landscape dominated by a broad extended basin.

Overall, we find that conformers **I** and **IV** merit further scrutiny because they exhibit the strongest deviations from MetaD-predicted behavior and reveal condition-sensitive kinetic trapping. Conformer **I** unexpectedly becomes immobilized in H^L^, while **IV** remains rigidly compact across all environments. These anomalous dynamical signatures indicate that specific electrostatic and salt-bridge interactions may be reshaping their local landscapes, prompting more focused structural and energetic analyses.

### How Electrostatics Modulate the Stability of the Canonical CaM Architecture

Conformer **I** corresponds to the 3CLN structure which is the canonical dumbbell-shaped conformation that has served for decades as the defining structural archetype of CaM.^9,54^ This geometry underpins innumerable mechanistic models of CaM function, from descriptions of calcium sensing to target recognition, allosteric communication, and linker flexibility. Because 3CLN is so widely regarded as the “reference” state of CaM, it is crucial to establish whether this conformation is genuinely stable across physiologically relevant ionic and loading conditions, or whether it is a crystallographically selected state that becomes destabilized in solution. In our cMD simulations, **I** exhibits condition-dependent drift and even kinetic trapping in low salt, behaviors at odds with its canonical status (**Figure 4**). These observations motivate a more detailed mechanistic examination of the electrostatic environment and ion–protein interactions that modulate the stability of this historically central CaM conformation.

We first probe the electrostatic environment surrounding conformer **I** by calculating the electrostatic potential along its surface using the Adaptive Poisson–Boltzmann Solver (APBS) tool,^55^ enabling visualization of the local electrostatic landscape. Surfaces are calculated with standard biomolecular and solvent dielectric constants (2 and 78.5, respectively), probe radius of 1.4 Å and with ionic compositions matched to each simulation condition of **Table 1**. Consistent with CaM’s strongly negative net charge at physiological pH (−23 for *apo*, −15 for *holo*), the protein surface is dominated by negative potential as shown in **Figure 5A**. However, the extent and spatial distribution of these negative regions depend strongly on ionic strength. We find that physiological salt partially neutralizes the electrostatic landscape, producing neutral patches in *apo* CaM and even small positive regions in the *holo* form mostly around regions of Ca^2+^ coordination. Low-salt conditions, in contrast, yield a uniformly negative surface that is expected to attract and retain counterions (K^+^) more strongly.

**Figure 5.**
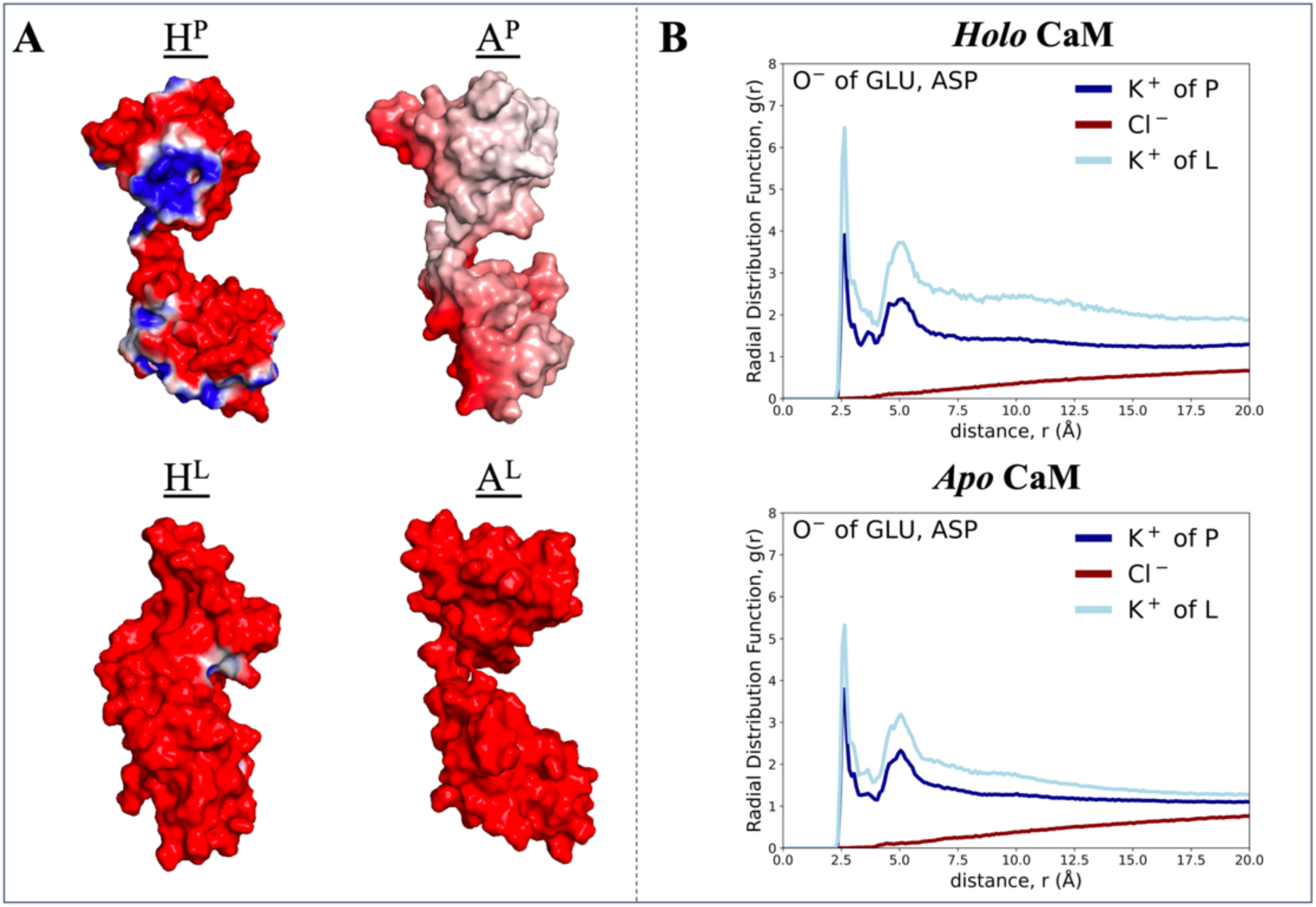
**(A)** Electrostatic isocontours of conformer **I** drawn at ± 1 *k*_B_*T/e*. Blue, red, and white regions represent positive, negative, and neutral electrostatic potentials, respectively, along the surface of the protein. **(B)** Radial distribution function, *g*(*r*), calculated between the C_β_ atoms of negatively charged residues and the surrounding K^+^ and Cl^−^ ions for the 200 ns-long simulations started from conformer I.

To quantify this ion association directly, we computed radial distribution functions (RDFs) between negatively charged side chains and surrounding ions for all simulation conditions (**Figure 5B**; for broader comparison see **Figure S5**). Across all conformers, a first coordination shell of K^+^ ions at 2–4 Å and a second one at 4–6 Å are consistently observed. Strikingly, more K^+^ ions approach the CaM surface under low-salt conditions, even though the number of K^+^ ions present in the physiological systems is significantly larger. This effect is robust and observed regardless of Ca^2+^ loading. The explanation becomes clear when examining Cl^−^ ion distributions, which under physiological salt form a diffuse co-ion layer that competes with K^+^ for proximity to the protein surface, weakening the electrostatic focusing of counterions. In low salt, however, the absence of co-ions leads to counterion condensation, producing tighter K^+^ clustering around acidic residues.

These ion-distribution trends are consistent with the electrostatic potential maps of **Figure 5A** such that physiological salt reduces and redistributes surface charge, while low salt enhances large contiguous negative patches that recruit more K^+^ ions. Such counterion accumulation can modulate conformational dynamics in two opposing ways: **(i)** by stabilizing compact electrostatically frustrated arrangements through local charge compensation, or **(ii)** by acting as “electrostatic lubricants” that weaken salt bridges and interdomain contacts, thereby enabling conformational rearrangements.

For **I**, the data suggest that both effects operate depending on ionic strength. Under physiological salt, weaker ion clustering and partial surface neutralization promote broader conformational mobility, consistent with the more diffuse sampling in H^P^ and especially in A^P^ (**Figure 4**). In the latter, the absence of Ca^2+^-mediated intradomain stabilization allows the two lobes to reorient more freely around the linker, yielding the widest coverage observed starting from the canonical dumbbell geometry. In fact, this condition even opens a route for this conformer to escape to the compact states exemplified by the 1PRW crystal structure. The conformational flip between 3CLN and 1PRW has been scrutinized in our earlier work where turning off the charge in E31 located in one of the EF-hand motifs enabled the compaction but not the rotation.^31^ Under low salt, however, dense K^+^ association reinforces local negative patches and enhances electrostatic steering, which, when combined with rigid Ca^2+^-loaded EF-hands in H^L^, promotes kinetic trapping of **I** in the shallow pockets of its landscape. In *apo* low-salt conditions (A^L^), the absence of EF-hand stabilization reduces this effect, and the conformer samples a somewhat broader region compared to H^L^.

Taken together, these analyses demonstrate that the stability of the canonical 3CLN conformation is not intrinsic but emergent, arising from a subtle balance among Ca^2+^-dependent intradomain rigidity, the redistribution of surface potential by ionic screening, and the local clustering of counterions at low ionic strength.

### How Salt-Bridge Networks Shape the Stability of the Compact CaM Architecture

Although counterion condensation introduces two competing effects, **(i)** stabilization of compact charge-frustrated regions and **(ii)** electrostatic lubrication that weakens salt bridges, their relative influence differs across conformers. Thus, while the same electrostatic processes operate across all conformers (**Figure S5**), the balance between stabilizing versus lubricating depends on the underlying geometry and packing of each state, which is reflected in the shifts in their RDF profiles. In the canonical dumbbell (3CLN), lubrication dominates, producing broad dispersion under physiological salt and kinetic trapping under H^L^. Similarly, the *cis*-extended **II** also lacks a compact interdomain interface minimizing the stabilizing effects, and its behavior across conditions is similar to conformer **I**. In contrast, the *cis*-compact **III** sits on a marginal ridge of all PMFs, where counterions preferentially weaken rather than reinforce interdomain contacts, facilitating its migration into neighboring basins across all conditions.

In contrast, the *trans*-compact CaM architecture, represented in our simulations by **IV** that remains compact across environments, possesses a pre-organized interdomain salt-bridge network that creates a stabilizing electrostatic clamp. This geometry uniquely benefits from the stabilizing effect of counterion condensation, as displayed by the largest K^+^ recruitment on the surface in both *apo* and *holo* forms, particularly in the low ionic strength conditions (**Figure S5**). To dissect the molecular basis of this rigidity, we quantified salt-bridge occupancies across conditions (**Table S1**) and traced the time evolution of interdomain charge–charge interactions for persistent salt bridges specific to this compact state (**Figure 6A**).

**Figure 6.**
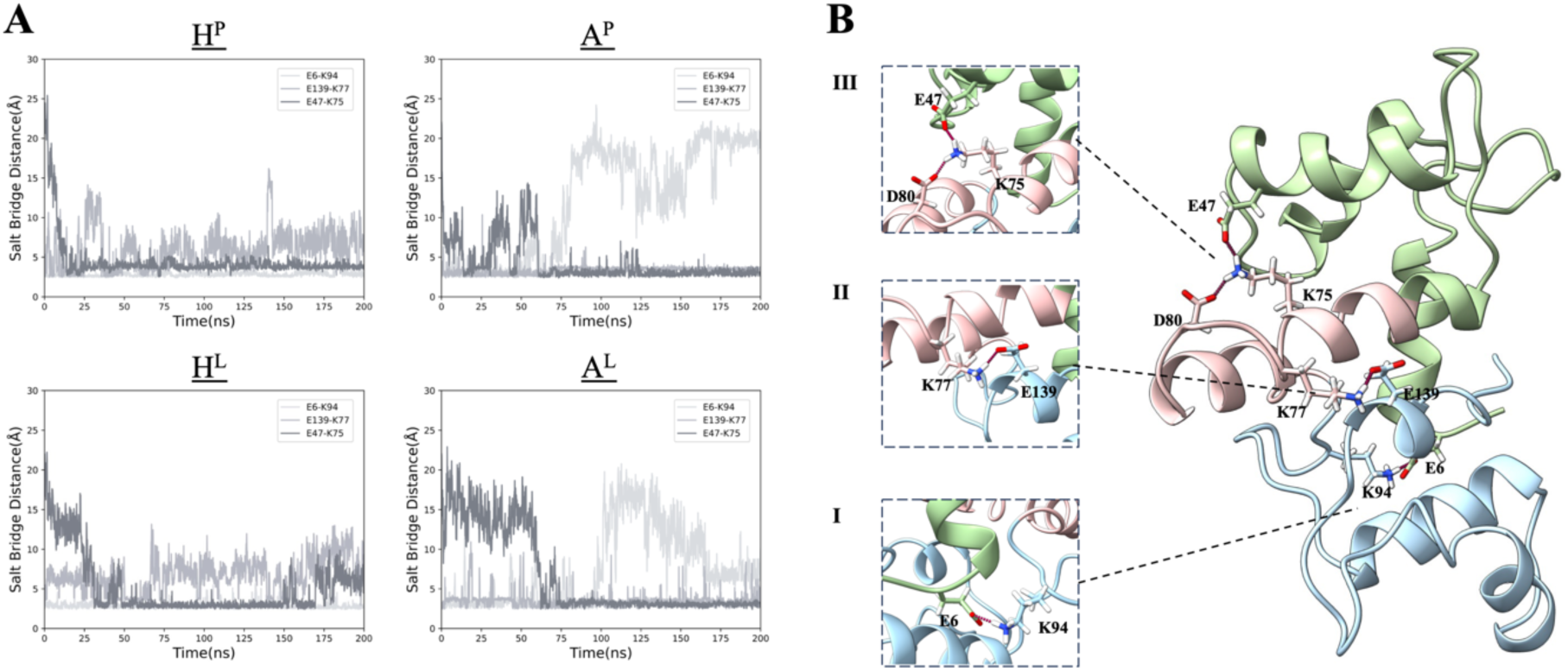
**(A)** Changes in unique salt bridge interactions of **IV** throughout the cMD trajectories for all systems, and **(B)** structural representations show salt bridge interactions identified in **IV** (snapshot selected from A^P^ simulation).

Three key electrostatic contacts underpin the stability of this compact architecture. First, the long-range salt bridge between E6 (N-lobe) and K94 (C-lobe) draws the two lobes inward and bends the helical linker (**Figure 6B-I**). This bridge is strong and persistent throughout the 200 ns simulations in both *holo* systems (H^P^, H^L^), the distance fluctuating between 3.2–5 Å (**Figure 5A**). It weakens earlier in the A^P^ trajectory and becomes intermittently labile in A^L^, reflecting the absence of Ca^2+^-stabilized EF-hand geometry. Second, the K77–E139 interaction couples the central linker to the C-lobe (**Figure 6B-II**). Though this bridge alternates between formed and broken states in H^P^ and H^L^, it remains remarkably stable in *apo* environments (A^P^ and A^L^), consistently residing near 3.2 Å. Third, the E47–K75 salt bridge (**Figure 6B-III**) forms only after the first two interactions have been established. In *apo* systems, it stabilizes after ~60 ns. In *holo* systems, it is established earlier on (<30 ns in these cMD simulations) and in H^L^ it becomes intermittently unstable after 170 ns, cycling between bound and unbound states.

These three bridges cooperatively contract the linker region and draw the lobes together. The cumulative effect is a multivalent interdomain clamp that restricts rotational freedom and enforces a compact geometry even under conditions where the free-energy landscape does not strongly favor it. The persistence of this compact state across ionic strengths and Ca^2+^-loading regimes therefore arises not from global landscape features but from the robustness of these salt-bridge networks, which act as internal stabilizers against both electrostatic lubrication and energetic flattening.

Together, these results show that compact CaM architectures benefit uniquely from counterion-modulated stabilization. Whereas extended or marginally compact states respond to ionic screening primarily through enhanced flexibility and basin-to-basin drift, the compact architecture is reinforced by a pre-existing interdomain electrostatic scaffold that remains partially intact across salt concentrations and Ca^2+^-loading regimes. In the A^P^ system in particular, ionic “lubrication” is insufficient to release the compact state: the same screening that softens long-range interactions leaves the short-range, multivalent E6–K94/K77–E139/E47–K75 clamp largely intact, so that concerted disruption of all contacts remains too rare to permit escape from this kinetically trapped configuration on the 200-ns time scale of the simulations. Although screening weakens individual interactions, it seldom breaks the entire network simultaneously. Since large-scale interdomain rearrangements require barrier-crossing events that cMD is unlikely to sample on these time scales, the compact state persists as the most kinetically rigid species in our simulations, even in conditions such as A^P^ and A^L^ where MetaD reveals no corresponding thermodynamic minimum, reflecting kinetic trapping enforced by the robust interdomain electrostatic network.

### How Cellular Context Selects CaM Conformations

CaM acts as a central interpreter of calcium signals across compartments that differ dramatically in ionic composition, calcium dynamics, and membrane architecture. Although it is primarily an intracellular protein found in the cytosol and nucleus,^13^ CaM also interacts with proteins in various calcium-regulating compartments^56^ (**Figure 7A**) where it not only regulates calcium levels but also plays a crucial role in calcium signaling by undergoing conformational changes that enable interactions with these proteins within organelles.^13^ This ability to regulate both calcium homeostasis and signaling demonstrates how CaM dynamically adjusts its structure to accommodate different binding partners. Its diverse localization, combined with its structural adaptability, allows it to function as a versatile regulator of calcium-dependent processes. This context-dependence makes CaM an ideal system to connect the MetaD-derived conformational landscapes to real biological function.

**Figure 7.**
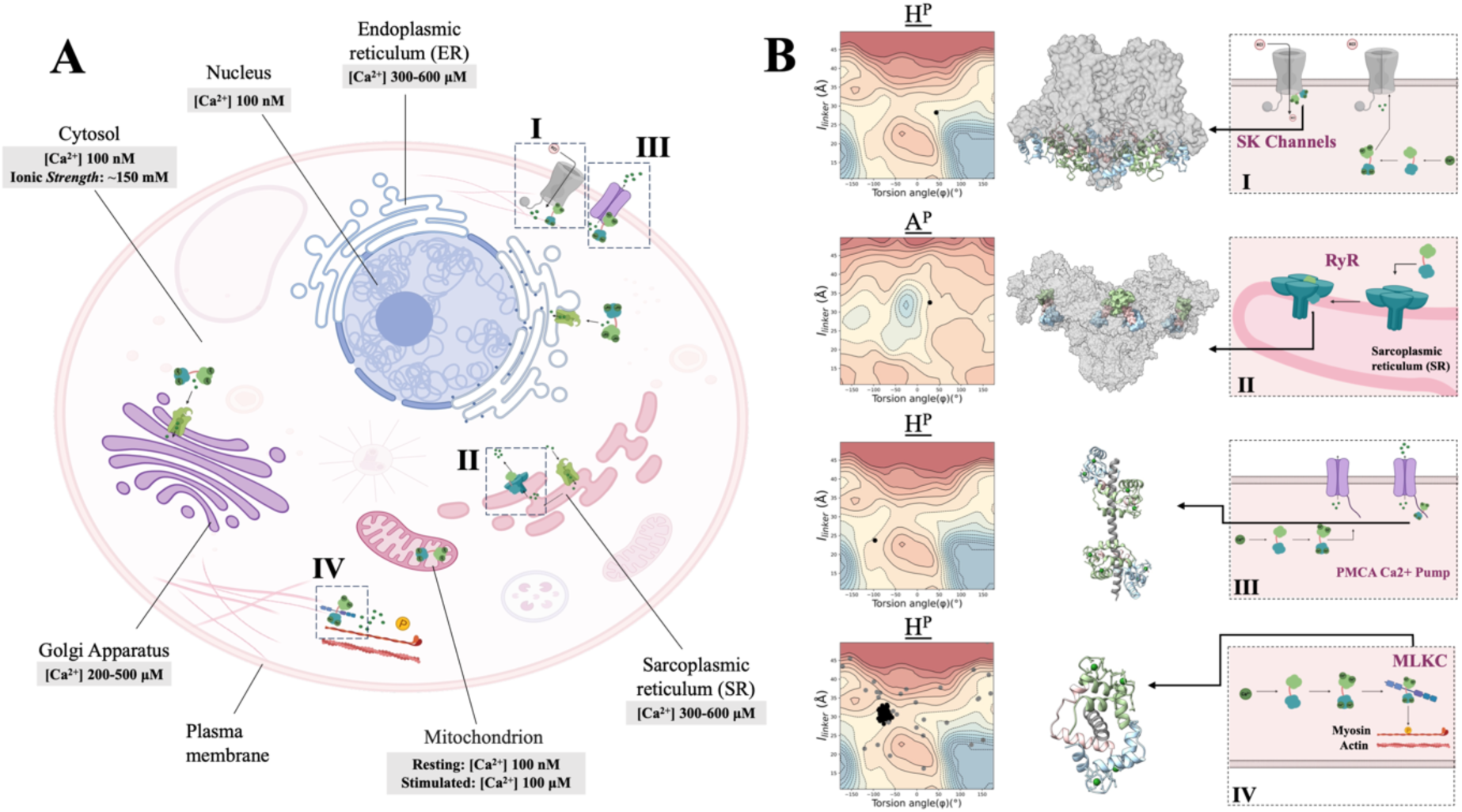
**(A)** A diagram of a eukaryotic cell shows the main organelles, typical calcium concentration distribution across compartments, and proteins located in different parts of the cell whose functions are regulated by CaM; **(B)** comparative conformational landscapes of CaM from MetaD simulations, overlaid with conformations of CaM from experimental structures with binding partners. Additionally, schematic illustrations depict how CaM interacts with these targets. CaM binding is shown for SK channel located in the plasma membrane (PDB ID: 6CNN) **(*i*)**, the RyR at the SR (PDB ID: 6JI8) **(*ii*)**, PMCA located in the plasma membrane (PDB ID: 4AQR) **(*iii*)** and MLCK in the cytoplasm (PDB IDs: 2K0F shown in black and 1MUX shown in gray) **(*iv*).**

To probe the functional significance of the conformational states sampled in our simulations, we projected experimental CaM–protein complex structures from multiple organelles onto our (*l*_linker_, *φ*) space (**Figure 7B**). We find that a coherent picture emerges whereby target-bound CaM overwhelmingly adopts *trans*-extended geometries, even when the target free, Ca^2+^-loaded protein favors a *cis*-compact state thermodynamically. This demonstrates that target engagement reshapes the CaM free-energy landscape, selecting conformations that are not the lowest-energy states in solution but become functionally stabilized through protein–protein interactions.

We first exemplify the conformational behavior of CaM in the plasma membrane. Here, CaM regulates both ion channels and Ca^2+^ extrusion machinery. Small-conductance calcium-activated potassium (SK) channels are membrane proteins that facilitate K^+^ flow across the cell membrane^57^ (**Figure 7A, i**). They are activated by intracellular Ca^2+^ but remain voltage-independent. Each SK channel tetramer associates constitutively with four CaM molecules,^58^ which act as built-in Ca^2+^ sensors. Upon Ca^2+^ binding, *holo* CaM fits into the SK channel cavity (**Figure 7B, i**). This interaction induces conformational changes that open the pore, allowing K^+^ ions to pass through.^57,58^ Without CaM, SK channels become inactive as they cannot respond to intracellular Ca^2+^ concentration changes. Cryo-EM structures show that *holo* CaM adopts a *cis*-extended geometry when bound to SK channels, precisely the region sampled in the shallow halo surrounding the H^P^ basin in our MetaD surface (**Figure 7B, i**). This conformation places both lobes in a configuration that can transmit Ca^2+^-dependent structural changes directly to the channel gating ring, providing a mechanistic explanation for how Ca^2+^ triggers pore opening.

On the sarcoplasmic reticulum (SR) ryanodine receptors (RyRs) are intracellular channels that mediate Ca^2+^ release into the cytoplasm^59^ (**Figure 7A, ii**). RyRs integrate both Ca^2+^-dependent and Ca^2+^-independent modes of CaM regulation.^60^ *apo* CaM suppresses RyR activity to prevent excessive Ca^2+^ leakage, while calcium-bound CaM fine-tunes RyR function during excitation-contraction coupling for controlled release.^61^ Cryo-EM structure of RyR bound to *apo*-CaM reveals that the N-lobe of CaM is positioned in the upper part of a cleft in RyR’s helical domain, while the C-lobe rests at the bottom, near the handle and central domains (**Figure 7B, ii**). When projected onto our MetaD landscapes, this structure aligns with the *cis*-extended minimum of the A^P^ surface, indicating that *apo* CaM binds RyR in a conformation that is naturally stable under physiological salt. This reinforces the established model in which *apo* CaM suppresses RyR leak by maintaining a tensioned, extended geometry stabilizing a closed-channel.

In the plasma membrane, CaM also interacts with Ca^2+^-ATPase (PMCA), a transport protein that moves Ca^2+^ from the cytosol to the extracellular space, maintaining low intracellular Ca^2+^ levels^62,63^ (**Figure 7A, iii**). When Ca^2+^ levels rise, *holo* CaM undergoes a conformational change and wraps around the C-terminal binding domain of PMCA, activating the pump and increasing Ca^2+^ efflux from the cell.^64^ Similar to SK, in its complex with PMCA, *holo* CaM adopts an arrangement that wraps around an 18-1 regulatory peptide (**Figure 7B, iii**). This binding-induced compaction corresponds to a region of the (*l*_linker_, *φ*) landscape that is accessible but not preferred by free *holo* CaM, consistent with the idea that PMCA binding stabilizes a structure that is otherwise only marginally sampled in the unbound ensemble, in essence leading to a redistribution of the conformational states.

In the cytoplasm, one well-known interaction partner of CaM is myosin light chain kinase (MLCK), a cytoplasmic enzyme associated with stress fibers and the cleavage furrow during cell division^65,66^ (**Figure 7A, iv**). It phosphorylates the regulatory light chain of myosin II, activating myosin for muscle contraction and other actomyosin-dependent processes.^65^ MLCK activity is significantly enhanced by CaM binding, whereas in its absence, the kinase remains largely inactive, impacting cellular signaling.^65^ Ca^2+^-loaded CaM activates MLCK by wrapping around a helical target peptide. In the NMR ensemble of the complex (PDB ID: 2K0F), CaM lies near the *cis*-extended region (**Figure 7B, iv**), with its C-lobe engaging the peptide and the N-lobe clamping around the target.^10^ This architecture lies well outside the deep *trans*-compact well of the H^P^ surface. Thus, peptide binding must reshape the CaM energy landscape, overriding the intrinsic *trans*-compact preference of calcium-loaded CaM and stabilizing an elongated geometry necessary for MLCK activation. The agreement of experimental structures with the accessible but non-dominant MetaD regions supports a model in which CaM’s intrinsic landscapes set the repertoire of possible regulatory states, while binding partners select and stabilize specific ones for function.

In contrast, overlaying the 30 conformers in the 1MUX NMR ensemble, having a structural and functional mimic of the CaM-binding region of MLCK,^67^ onto the MetaD-derived landscape (**Figure 7B, iv**) shows that the *holo* CaM structures in this complex populate even further extended conformations with *l*_linker_ > 35 Å. Rather than occupying the deep *trans*-compact minimum that characterizes free *holo* CaM, the ensemble represents extended states that are only weakly sampled in our simulations. By occupying MLCK-recognition pockets, W-7 forcibly prevents lobe–lobe collapse and stabilizes extended CaM geometries, effectively overriding the intrinsic *trans*-compact preference of Ca^2+^-loaded CaM. Thus, the 1MUX ensemble provides a clear structural demonstration that MLCK-like target engagement can recruit *holo* CaM into elongated states located outside its intrinsic energetic minimum, consistent with the extended regions partially accessed in our MetaD surface.

The contrast between the 1MUX and 2K0F ensembles highlights how different target peptides sculpt distinct regions of the CaM energy landscape. In 1MUX, the flexible W-7 peptide provides only weak geometric constraints and allows *holo* CaM to populate a wide continuum of elongated states, spanning both *cis*- and *trans*-extended geometries. In 2K0F, however, the CaM-binding partner is a preorganized amphipathic helix with a fixed hydrophobic register, enforcing a single interdomain orientation and collapsing the ensemble onto a narrow band in the *cis*-region. Thus, the breadth or specificity of CaM’s extended conformational states is determined not only by its intrinsic landscape but by the structural rigidity and anchoring pattern of the bound target peptide.

Taken together, the cross-compartment examples highlight unifying trends for CaM conformational selection: First, *apo* CaM functions predominantly in extended geometries, consistent with our A^P^ and A^L^ free-energy surfaces. This underlies its regulatory roles in preventing aberrant Ca^2+^ release (RyR). Second, Ca^2+^ loading introduces a deep *trans*-compact basin, which is biologically relevant for peptide-free CaM but is rarely used directly for target binding. Instead, Ca^2+^-loaded CaM typically transitions into less compact regions upon engaging partners like SK channels, PMCA, and MLCK. This engagement may also help release the deeply stabilizing linker – lobe salt bridges of CaM (**Figure 6**). Third, target binding consistently recruits CaM into higher-lying regions of the MetaD landscape, demonstrating that functional conformations are often not the global thermodynamic minima of the isolated protein. Instead, protein–protein contacts stabilize geometries that are subdominant or transient in solution.

In sum, ligand binding reshapes CaM’s charge distribution and steric constraints in ways that disrupt compact-state salt-bridge stabilization and redirect the protein into specific elongated conformations drawn from its intrinsic solution-phase landscape. Interpreting simulations with experimental structures illustrates clearly how CaM’s environment, i.e., ionic strength, calcium loading, organellar context, and binding partner, collectively reshapes its conformational ensemble. The result is a dynamic, context-sensitive regulator capable of executing distinct tasks across the cell.

## Conclusions and Future Perspectives

CaM’s versatility as a calcium sensor emerges from a finely tuned interplay between Ca^2+^ binding, ionic strength, and intrinsic electrostatics, all of which reshape a structured yet adaptable conformational landscape. By combining well-tempered MetaD with classical MD simulations, and interpreting our findings with experimentally determined structures, we show that CaM’s conformational ensemble is not merely flexible but systematically regulated by environmental conditions and requirements for engagement with specific targets.

Under physiological ionic strength (**Figure 2**), Ca^2+^ loading drives CaM into a deeply stabilized *trans*-compact basin, reflecting a rigidified EF-hand architecture supported by an interdomain salt-bridge network that restricts linker mobility. *apo* CaM, in contrast, favors a broad *cis*-extended ensemble with enhanced rotational freedom of the lobes. Reducing the ionic strength (**Figure 3**) greatly flattens the free-energy surfaces for both states, lowering barriers and enabling facile interconversion. Yet, this flattening simultaneously promotes spurious electrostatic trapping, as strong K^+^ condensation around acidic residues stabilizes local minima that are bypassed in MetaD but kinetically persistent in unbiased simulations (**Figure 4**).

Comparing MetaD and cMD results reveals that while MetaD exposes CaM’s thermodynamic possibilities, cMD reveals which of those states are kinetically stable. Many higher-lying MetaD basins dissipate under unbiased dynamics, while one structure, the *trans*-compact conformer, remains strikingly immobile across all conditions due to a network of salt-bridges (**Figure 6**). This rigid architecture underscores how short-range electrostatics can override broader energetic trends.

A coherent picture emerges when the conformational landscapes of free CaM are considered alongside ligand-bound structures across cellular compartments. In fact, overlaying experimentally observed CaM–target complexes onto our landscapes highlights that functional CaM conformations often lie outside thermodynamic minima (**Figure 7**). In solution, Ca^2+^-loaded CaM is governed by a deep *trans*-compact basin stabilized by an interdomain salt-bridge network (**Figure 6**), whereas *apo* CaM favors a broad *cis*-extended ensemble with few geometric constraints. Ionic strength modulates the ease with which CaM moves between these states by reshaping counterion condensation and the stability of interdomain contacts (**Figure 5**). Target engagement shifts this balance not by altering ionic composition but by reorganizing CaM’s charge distribution and steric geometry. Binding partners, whether flexible antagonists for MLCK such as W-7 in 1MUX or highly structured regulatory helices as in 2K0F, or membrane-associated peptides from SK channels and PMCA, bury acidic pockets, generate new hydrophobic anchors, and sterically block formation of the compact-state salt-bridge clamp. As a result, the multivalent electrostatic scaffold that stabilizes the *trans*-compact minimum is dismantled, and CaM is redirected into elongated conformations that match the functional geometry required by each target. Thus, ligand binding selectively stabilizes higher-lying but readily accessible regions of CaM’s intrinsic energy landscape, allowing the protein to adopt distinct regulatory states across different cellular contexts.

Together, these results provide an integrated view of CaM as a protein whose conformational preferences are not fixed but emergent outcomes of electrostatics, ion composition, and target-induced remodeling. They also clarify long-standing discrepancies between crystallographicly determined states, NMR ensembles, and solution-phase biophysical measurements, showing how each corresponds to a different region of the underlying multidimensional landscape.

We note that while the rotation–linker distance CV pair offers clear interpretability and successfully recapitulates known conformers, it remains a reduced projection of CaM’s high-dimensional dynamics. Exploring alternative collective variables in future studies may reveal additional slow modes or refine the boundaries between basins. Future work could also focus on identifying transition pathways between conformers, incorporating experimental restraints from FRET or MS, or simulating explicit target-binding events to map how partner engagement deforms the energy landscape. Ultimately, the new conformers and mechanistic insights identified here provide a foundation for predictive modeling of CaM regulation and for designing synthetic peptides or mutations that tune CaM’s conformational equilibria.

## Methods

### System Preparation

To investigate how the absence or presence of Ca^2+^ ions and changes in salt concentration influence CaM’s conformations, we performed classical MD (cMD) and well-tempered MetaD simulations (labelled simply MetaD in the rest of the manuscript) under each condition (**Table 1**). The calcium-loaded, extended structure of CaM serves as the canonical form of CaM in the literature; thus, we used the 3CLN structure (**Figure 1**) as the starting point for system preparation. We labelled the systems as H^P^ (calcium-loaded *holo* CaM at physiological salt concentration), H^L^ (*holo* CaM in low salt), A^P^ (Ca^2+^-free *apo* CaM at physiological salt concentration) and A^L^ (*apo* CaM at low salt), respectively. To simulate *apo* conditions in these initial systems, Ca^2+^ ions were removed from the 3CLN structure. By using the solvate plug-in of VMD,^47^ we solvated the protein structures in a rectangular water box with a minimum distance of 10 Å between the protein and the nearest edge. We added K^+^ ions to neutralize charges in low salt simulations, with additional K^+^ and Cl^−^ ions to mimic physiological ion concentration (see **Table 1**).

### cMD Simulations

We performed duplicate 1*μ*s long cMD simulations under each condition by using the NAMD software package,^48^ with the CHARMM36 force-field to model the protein and the TIP3P model for water molecules.^49^ We utilized VMD for preprocessing of structures, such as protein structure file (PSF) generation, solvation (constructing water-box), ionization and visualization of MD trajectories.^47^ We employed particle mesh Ewald method for calculating long-range electrostatic interactions with a cutoff of 12 Å with a switching distance of 10 Å. We applied the RATTLE algorithm to constrain bonds, enabling a 2 fs timestep with Verlet integration. We used Langevin piston for pressure control at 1 atm, and we controlled the temperature at 310 K by the Langevin thermostat. We minimized each system for 10000 steps, and performed equilibrium simulations in the *NPT* ensemble. We stored the trajectories every 100 ps.

We used the first frame of each trajectory as a reference for root-mean-square deviation (RMSD) calculations. We performed radial distribution function, *g*(*r*), calculations in VMD with the periodic boundary conditions taken into account and the distances binned in 0.1 Å increments.^47^ For these calculations, we selected K^+^ and Cl^−^ ions as the first reference group components, and the side chains of negatively charged residues (GLU or ASP) as the second reference group. We carried out trajectory analyses and RMSD calculations with the ProDy package^50^ within the Python programming environment. We made all in-house scripts used in this study available on GitHub.

### Well-Tempered Metadynamics (MetaD) Simulations

Two geometry-related CVs were defined to efficiently explore the conformational landscape of CaM (**Figure 1B**). To represent the switch between extended and compact conformations of CaM, following the methodology we introduced in prior studies,^31^ one CV is the linker end-to-end distance (*l_linker_*) defining the distance between the C_α_ atoms of residues 69 and 91 marking the end points of the linker region. The second CV is the coarse-grained torsion angle (*φ*) describing the positioning of the two lobes relative to each other. This torsion angle is defined using, in order, the center of mass of the N-terminal domain, the C_α_ atoms at the beginning and end of the helical linker region, and the center of mass of the C-terminal domain.

We selected snapshots from MD simulation trajectories for each condition as starter structures for MetaD simulations. During this selection phase, we focused on the RMSD and the defined CVs. We selected the most stable conformer under each condition specifically by targeting regions with minimal changes in RMSD (**Figure S1**). Additionally, (*l*_linker_, *φ*) plots (**Figure S2**) were crucial in determining the conformational range sampled in these trajectories. After selecting the snapshots, we removed the water molecules, K^+^ and Cl^−^ ions and followed the same system preparation procedure as in the cMD simulations: place the structures in a water box and add enough ions to achieve salt concentrations corresponding to the conditions under which the MetaD simulations are to be carried out.

We minimized each system for 10000 steps, followed by a 100 ns equilibrium simulation to reach a stable local minimum. This approach ensured that the starting configurations for the MetaD simulations began at well-defined minima, allowing Gaussian hills to be deposited effectively for enhanced sampling. We then started MetaD simulations using the final atomic coordinates and velocities. We defined the upper and lower limits of *l*_linker_ as 10–50 Å, divided into 1.5 Å grid intervals, while we divided *φ* into 10° grids spanning the range [-π,π]. The height and width of the Gaussian were 0.2 kcal mol^-1^ and 1.0 Å respectively. The bias factor for well-tempering was set to 6, and hills were deposited every 500 steps. All MetaD simulations were carried out for 600 ns.

### cMD Simulations of four MetaD sampled conformers under different conditions

We selected four conformers representing the compact/extended structures where the two lobes face the same side (*cis* arrangement) or opposite sides (*trans* arrangement) of the linker from the MetaD simulations for further tests of their stability under the prevailing conditions. Selection and labeling of these structures are detailed in the Results and Discussion section. We subjected each of these structures to additional MD simulations under the conditions of H^P^, A^P^, H^L^ and A^L^. After selecting the conformers, we removed water molecules and salts from these snapshots. We carried out 10000-step minimization followed by 200 ns run for each structure, using the same MD simulation protocol as the original cMD simulations.

## Supporting Information

Five figures (S1-S5) and one table (S1).

## Acknowledgments

The numerical calculations reported in this paper were partially performed at the TÜBİTAK ULAKBİM High Performance and Grid Computing Center (TRUBA resources). This work was financially supported by TÜBİTAK project no. 122F149. We thank Dilara Çoban for contributions during the initial stages of this study.

## Author Contributions

B.T. performed the calculations and wrote the manuscript. S.H. contributed to the initial computational analyses. A.R.A. contributed to the design of the study, co-supervised the project and participated in discussions and manuscript preparation. C.A. secured funding, conceived and designed the work, supervised the project, and wrote the manuscript.

## Data and Code Availability

Original code for producing the results for this study is deposited at GitHub at the following repository: https://github.com/midstlab/Tayhan_2026. All MD simulation trajectories produced in this work will be provided by the lead contact upon request.

## References

1 Berridge, M. J. et al. The versatility and universality of calcium signalling. Nature Reviews Molecular Cell Biology 1, 11–21 (2000). 10.1038/35036035

2 Raffaello, A., Mammucari, C., Gherardi, G. & Rizzuto, R. Calcium at the Center of Cell Signaling: Interplay between Endoplasmic Reticulum, Mitochondria, and Lysosomes. Trends in Biochemical Sciences 41, 1035–1049 (2016). 10.1016/j.tibs.2016.09.001

3 Banci, L. & Bertini, I. Metallomics and the cell: some definitions and general comments. Metal ions in life sciences 12, 1–13 (2013). 10.1007/978-94-007-5561-1_1

4 Kaufman, R. J. & Malhotra, J. D. Calcium trafficking integrates endoplasmic reticulum function with mitochondrial bioenergetics. Biochimica et Biophysica Acta (BBA) - Molecular Cell Research 1843, 2233–2239 (2014). 10.1016/j.bbamcr.2014.03.022

5 Dolman, N. & Tepikin, A. Calcium Gradients and the Golgi. Cell calcium 40, 505–512 (2006). 10.1016/j.ceca.2006.08.012

6 Duchen, M. R. Mitochondria and calcium: from cell signalling to cell death. Journal of Physiology 529, 57–68 (2000). 10.1111/j.1469-7793.2000.00057.x

7 Bagur, R. & Hajnóczky, G. Intracellular Ca2+ Sensing: Its Role in Calcium Homeostasis and Signaling. Molecular Cell 66, 780–788 (2017). 10.1016/j.molcel.2017.05.028

8 Carafoli, E., Santella, L., Branca, D. & Brini, M. Generation, Control, and Processing of Cellular Calcium Signals. Critical Reviews in Biochemistry and Molecular Biology 36, 107–260 (2001). 10.1080/20014091074183

9 Chin, D. & Means, A. R. Calmodulin: a prototypical calcium sensor. Trends in Cell Biology 10, 322–328 (2000). 10.1016/s0962-8924(00)01800-6

10 Gsponer, J. et al. A Coupled Equilibrium Shift Mechanism in Calmodulin-Mediated Signal Transduction. Structure 16, 736–746 (2008). 10.1016/j.str.2008.02.017

11 Babu, Y. S., Bugg, C. E. & Cook, W. J. Structure of calmodulin refined at 2.2 Å resolution. Journal of Molecular Biology 204, 191–204 (1988). 10.1016/0022-2836(88)90608-0

12 Nelson, M. & Chazin, W. Structures of EF-hand Ca(2+)-binding Proteins: Diversity in the Organization, Packing and Response to Ca2+ binding. Biometals 11, 297–318 (1998). 10.1023/a:1009253808876

13 Vetter, S. & Leclerc, E. Novel aspects of calmodulin target recognition and activation. European Journal of Biochemistry 270, 404–414 (2003). 10.1046/j.1432-1033.2003.03414.x

14 Negi, S., Aykut, A. O., Atilgan, A. R. & Atilgan, C. Calmodulin Readily Switches Conformation upon Protonating High pK(a) Acidic Residues. Journal of Physical Chemistry B 116, 7145–7153 (2012). 10.1021/jp3032995

15 Kuboniwa, H. et al. Solution structure of calcium-free calmodulin. Nature Structural Biology 2, 768–776 (1995). 10.1038/nsb0995-768

16 Villalobo, A., Ishida, H., Vogel, H. J. & Berchtold, M. W. Calmodulin as a protein linker and a regulator of adaptor/scaffold proteins. Biochimica et Biophysica Acta (BBA) - Molecular Cell Research 1865, 507–521 (2018). 10.1016/j.bbamcr.2017.12.004

17 Westerlund, A. M. & Delemotte, L. Effect of Ca2+ on the promiscuous target-protein binding of calmodulin. PLOS Computational Biology 14, e1006072 (2018). 10.1371/journal.pcbi.1006072

18 Fiorin, G., Pastore, A., Carloni, P. & Parrinello, M. Using Metadynamics to Understand the Mechanism of Calmodulin/Target Recognition at Atomic Detail. Biophysical Journal 91, 2768–2777 (2006). 10.1529/biophysj.106.086611

19 Grant, B. M. M., Enomoto, M., Ikura, M. & Marshall, C. B. A Non-Canonical Calmodulin Target Motif Comprising a Polybasic Region and Lipidated Terminal Residue Regulates Localization. International Journal of Molecular Sciences 21, 2751 (2020). 10.3390/ijms21082751

20 Swindells, M. & Ikura, M. Pre-formation of the semi-open conformation by the apo-calmodulin C-terminal domain and implications binding IQ-motifs. Nature Structural Biology 3, 501–504 (1996). 10.1038/nsb0696-501

21 Atilgan, A. R., Aykut, A. O. & Atilgan, C. Subtle pH differences trigger single residue motions for moderating conformations of calmodulin. Journal of Chemical Physics 135, 155102 (2011). 10.1063/1.3651807

22 Zhang, M. et al. Structural Basis for Calmodulin as a Dynamic Calcium Sensor. Structure 20, 911–923 (2012). 10.1016/j.str.2012.03.019

23 Brian D. Slaughter, Michael W. Allen, Jay R. Unruh, Ramona J. Bieber Urbauer, A. & Johnson, C. K. Single-Molecule Resonance Energy Transfer and Fluorescence Correlation Spectroscopy of Calmodulin in Solution†. Journal of Physical Chemistry B 108, 10388–10397 (2004). 10.1021/jp040098

24 Wyttenbach, T., Grabenauer, M., Thalassinos, K., Scrivens, J. H. & Bowers, M. T. The Effect of Calcium Ions and Peptide Ligands on the Relative Stabilities of the Calmodulin Dumbbell and Compact Structures. Journal of Physical Chemistry B 114, 437–447 (2009). 10.1021/jp906242

25 Berman, H. M. et al. The Protein Data Bank. Nucleic Acids Res 28, 235–242 (2000). 10.1093/nar/28.1.235

26 Fallon, J. L. & Quiocho, F. A. A Closed Compact Structure of Native Ca2+-Calmodulin. Structure 11, 1303–1307 (2003). 10.1016/j.str.2003.09.004

27 Jiang, J. et al. Site-specific modification of calmodulin Ca2+ affinity tunes the skeletal muscle ryanodine receptor activation profile. Biochemical Journal 432, 89–99 (2010). 10.1042/BJ20100505

28 Hoeflich, K. P. & Ikura, M. Calmodulin in Action: Diversity in Target Recognition and Activation Mechanisms. Cell 108, 739–742 (2002). 10.1016/S0092-8674(02)00682-7

29 Holt, C. et al. The arrhythmogenic N53I variant subtly changes the structure and dynamics in the calmodulin N-terminal domain, altering its interaction with the cardiac ryanodine receptor. Journal of Biological Chemistry 295, 7620–7634 (2020). 10.1074/jbc.RA120.013430

30 Brian D. Slaughter, Michael W. Allen, Jay R. Unruh, Ramona J. Bieber Urbauer, A. & Johnson*, C. K. Single-Molecule Resonance Energy Transfer and Fluorescence Correlation Spectroscopy of Calmodulin in Solution†. (June 11, 2004). 10.1021/jp040098

31 Aykut, A. O., Atilgan, A. R. & Atilgan, C. Designing Molecular Dynamics Simulations to Shift Populations of the Conformational States of Calmodulin. PLOS Computational Biology 9, e1003366 (2013). 10.1371/journal.pcbi.1003366

32 Abdizadeh, H., Atilgan, A. R. & Atilgan, C. Mechanisms by Which Salt Concentration Moderates the Dynamics of Human Serum Transferrin. Journal of Physical Chemistry B 121, 4778–4789 (2017). 10.1021/acs.jpcb.7b02380

33 Sensoy, O., Atilgan, A. R. & Atilgan, C. FbpA iron storage and release are governed by periplasmic microenvironments. Physical Chemistry Chemical Physics 19, 6064–6075 (2017). 10.1039/c6cp06961d

34 Bussi, G., Laio, A., Bussi, G. & Laio, A. Using metadynamics to explore complex free-energy landscapes. Nature Reviews Physics 2, 200–212 (2020). 10.1038/s42254-020-0153-0

35 Laio, A. & Parrinello, M. Escaping free-energy minima. Proceedings of the National Academy of Sciences of the United States of America 99, 12562–12566 (2002). 10.1073/pnas.202427399

36 Sugita, Y. & Okamoto, Y. Replica-exchange molecular dynamics method for protein folding. Chemical Physics Letters 314, 141–151 (1999). 10.1016/S0009-2614(99)01123-9

37 Torrie, G. M. & Valleau, J. P. Nonphysical sampling distributions in Monte Carlo free-energy estimation: Umbrella sampling. Journal of Computational Physics 23, 187–199 (1977). 10.1016/0021-9991(77)90121-8

38 Comer, J. et al. The Adaptive Biasing Force Method: Everything You Always Wanted To Know but Were Afraid To Ask. Journal of Physical Chemistry B 119, 1129–1151 (2014). 10.1021/jp506633

39 Laio, A., Gervasio, F. L., Laio, A. & Gervasio, F. L. Metadynamics: a method to simulate rare events and reconstruct the free energy in biophysics, chemistry and material science. Reports on Progress in Physics 71, 126601 (2008). 10.1088/0034-4885/71/12/126601

40 Barducci, A., Bussi, G. & Parrinello, M. Well-Tempered Metadynamics: A Smoothly Converging and Tunable Free-Energy Method. Physical Review Letters 100, 020603 (2008). 10.1103/PhysRevLett.100.020603

41 McCarty, J. & Parrinello, M. A variational conformational dynamics approach to the selection of collective variables in metadynamics. The Journal of Chemical Physics 147, 204109 (2017). 10.1063/1.4998598

42 Ray, D. & Parrinello, M. Kinetics from Metadynamics: Principles, Applications, and Outlook. Journal of Chemical Theory and Computation 19, 5649–5670 (2023). 10.1021/acs.jctc.3c00660

43 Pérez-Hernández, G., Paul, F., Giorgino, T., De Fabritiis, G. & Noé, F. Identification of slow molecular order parameters for Markov model construction. Journal of Chemical Physics 139, 015102 (2013). 10.1063/1.4811489

44 Maria-Solano, M. A., Serrano-Hervás, E., Romero-Rivera, A., Iglesias-Fernández, J. & Osuna, S. Role of conformational dynamics in the evolution of novel enzyme function. Chemical Communications 54, 6622–6634 (2018). 10.1039/c8cc02426j

45 Atilgan, A. & Atilgan, C. Computational strategies for protein conformational ensemble detection. Current Opinion in Structural Biology 72, 79–87 (2022).

46 McCoy, M. D., John Hamre, I., Klimov, D. K. & Jafri, M. S. Predicting Genetic Variation Severity Using Machine Learning to Interpret Molecular Simulations. Biophysical Journal 120, 189–204 (2020). 10.1016/j.bpj.2020.12.002

47 Humphrey, W., Dalke, A. & Schulten, K. VMD: visual molecular dynamics. Journal of Molecular Graph 14, 33–38 (1996). 10.1016/0263-7855(96)00018-5

48 Phillips, J. C. et al. Scalable molecular dynamics on CPU and GPU architectures with NAMD. The Journal of Chemical Physics 153, 044130 (2020). 10.1063/5.0014475

49 Best, R. B. et al. Optimization of the Additive CHARMM All-Atom Protein Force Field Targeting Improved Sampling of the Backbone ϕ, ψ and Side-Chain χ1 and χ2 Dihedral Angles. Journal of Chemical Theory and Computation 8, 3257–3273 (2012). 10.1021/ct300400x

50 Bakan, A., Meireles, L. M. & Bahar, I. ProDy: protein dynamics inferred from theory and experiments. Bioinformatics 27, 1575–1577 (2011). 10.1093/bioinformatics/btr168

51 Barbato, G., Ikura, M., Kay, L. E., Pastor, R. W. & Bax, A. Backbone dynamics of calmodulin studied by nitrogen-15 relaxation using inverse detected two-dimensional NMR spectroscopy: the central helix is flexible. Biochemistry 31, 5269–5278 (1992). 10.1021/bi00138a005

52 Linse, S., Helmersson, A. & Forsén, S. Calcium binding to calmodulin and its globular domains. Journal of Biological Chemistry 266, 8050–8054 (1991). 10.1016/S0021-9258(18)92938-8

53 Chou, J. J., Li, S. P., Klee, C. B. & Bax, A. Solution structure of Ca2+-calmodulin reveals flexible hand-like properties of its domains. Nature Structural Biology 8, 990–997 (2001). 10.1038/nsb1101-990

54 Finn, B. E. & Forsén, S. The evolving model of calmodulin structure,function and activation. Structure 3, 7–11 (1995). 10.1016/S0969-2126(01)00130-7

55 Jurrus, E. et al. Improvements to the APBS biomolecular solvation software suite. Protein Science 27, 112–128 (2018). 10.1002/pro.3280

56 Clapham, D. E. Calcium signaling. Cell 80, 259–268 (1995). 10.1016/0092-8674(95)90408-5

57 Lee, C.-H. & MacKinnon, R. Activation mechanism of a human SK-calmodulin channel complex elucidated by cryo-EM structures. Science 360, 508–513 (2018). 10.1126/science.aas9466

58 Gu, M. et al. Small-conductance Ca2+-activated K+ channels: insights into their roles in cardiovascular disease. Experimental & Molecular Medicine 50, 1–7 (2018). 10.1038/s12276-018-0043-z

59 Xu, L. & Meissner, G. Mechanism of Calmodulin Inhibition of Cardiac Sarcoplasmic Reticulum Ca2+ Release Channel (Ryanodine Receptor). Biophysical Journal 86, 797–804 (2004). 10.1016/S0006-3495(04)74155-7

60 Balshaw, D. M., Xu, L., Yamaguchi, N., Pasek, D. A. & Meissner, G. Calmodulin Binding and Inhibition of Cardiac Muscle Calcium Release Channel (Ryanodine Receptor). Journal of Biological Chemistry 276, 20144–20153 (2001). 10.1074/jbc.M010771200

61 Gong, D. et al. Modulation of cardiac ryanodine receptor 2 by calmodulin. Nature 572, 347–351 (2019). 10.1038/s41586-019-1377-y

62 Tidow, H. et al. A bimodular mechanism of calcium control in eukaryotes. Nature 491, 468–472 (2012). 10.1038/nature11539

63 Berrocal, M. & Mata, A. M. The Plasma Membrane Ca2+-ATPase, a Molecular Target for Tau-induced Cytosolic Calcium Dysregulation. Neuroscience 518, 112–118 (2023). 10.1016/j.neuroscience.2022.04.016

64 Juranic, N. et al. Calmodulin Wraps around Its Binding Domain in the Plasma Membrane Ca2+ Pump Anchored by a Novel 18-1 Motif. Journal of Biological Chemistry 285, 4015–4024 (2010). 10.1074/jbc.M109.060491

65 Poperechnaya, A., Varlamova, O., Lin, P.-J., Stull, J. T. & Bresnick, A. R. Localization and Activity of Myosin Light Chain Kinase Isoforms during the Cell Cycle. The Journal of Cell Biology 151, 697–708 (2000). 10.1083/jcb.151.3.697

66 Kamm, K. E. & Stull, J. T. Dedicated Myosin Light Chain Kinases with Diverse Cellular Functions *. Journal of Biological Chemistry 276, 4527–4530 (2001). 10.1074/jbc.R000028200

67 Osawa, M. et al. Solution structure of calmodulin-W-7 complex: The basis of diversity in molecular recognition. Journal of Molecular Biology 276, 165–176 (1998). 10.1006/jmbi.1997.1524

