## Supplementary information for "Functional Calmodulin States are Selected from an Electrostatically Tuned Free Energy Landscape"

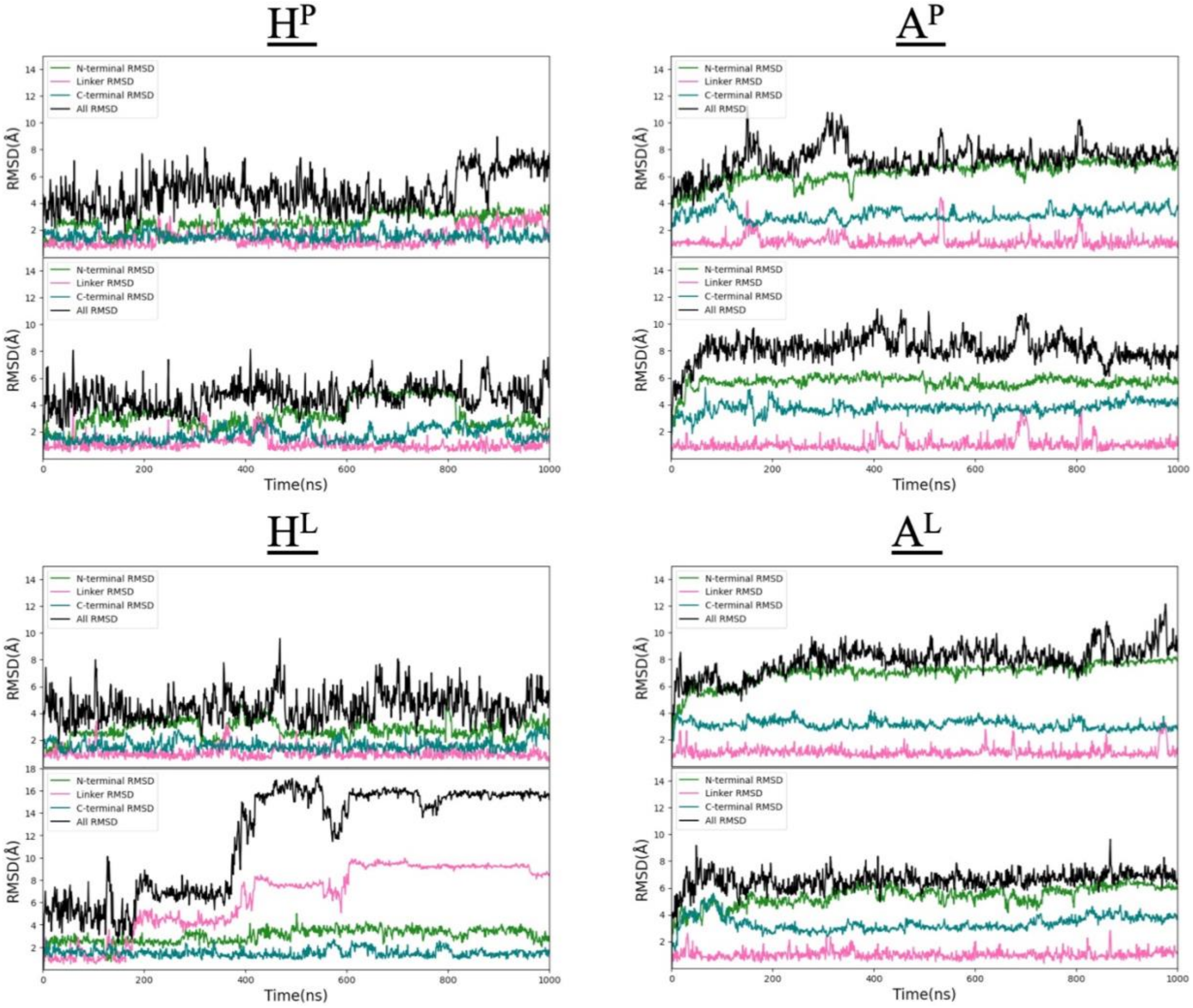


**Figure S1 |** RMSD graphs of two independent 1 μs long simulations of CaM in different environmental conditions; $H^P$, $A^P$, $H^L$ and $A^L$. N-terminal domain, C-terminal domain, flexible linker region, and overall RMSD are depicted in green, blue, pink, and black, respectively. The colors of the regions correspond to the coloring displayed in **Figure 1**.

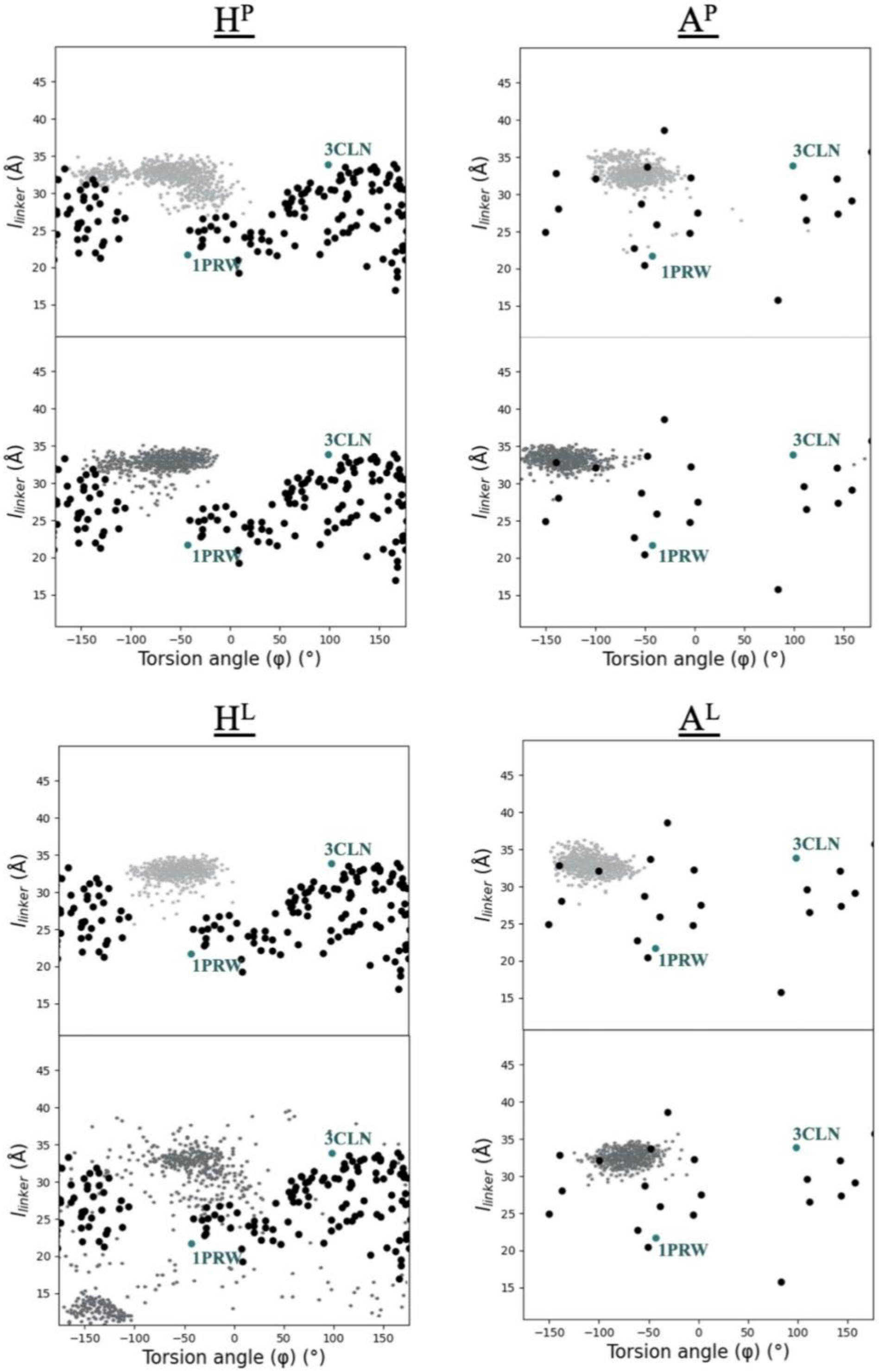


**Figure S2** | Torsion angle ($\varphi$) – linker end-to-end distance ($l_{\text{linker}}$) plots from duplicate cMD trajectories (light gray/dark gray small dots) under the four environmental conditions studied, all started from the 3CLN initial structure. For reference, experimental structures are also overlaid with the larger dots: Black ones represent NMR structures determined for *holo* (2K0E; 160 conformers) or *apo* CaM (6Y95; 20 conformers); in teal color are two labelled crystal structures.

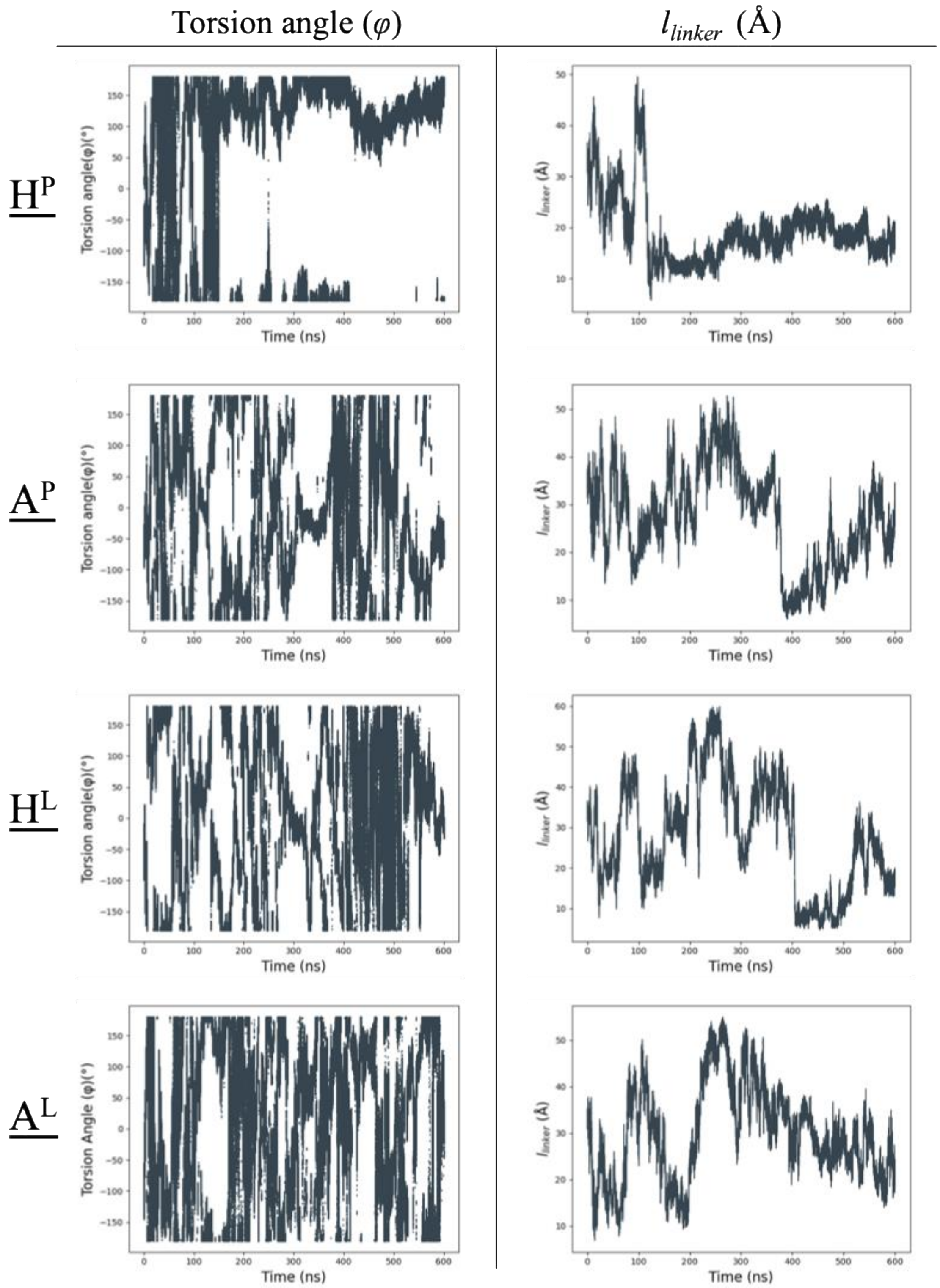


**Figure S3** │ wt-MetaD trajectories for the two selected CVs: the left panel shows changes in the torsional angle (°), while the right panel displays the linker end-to-end distance (Å) over time across all systems

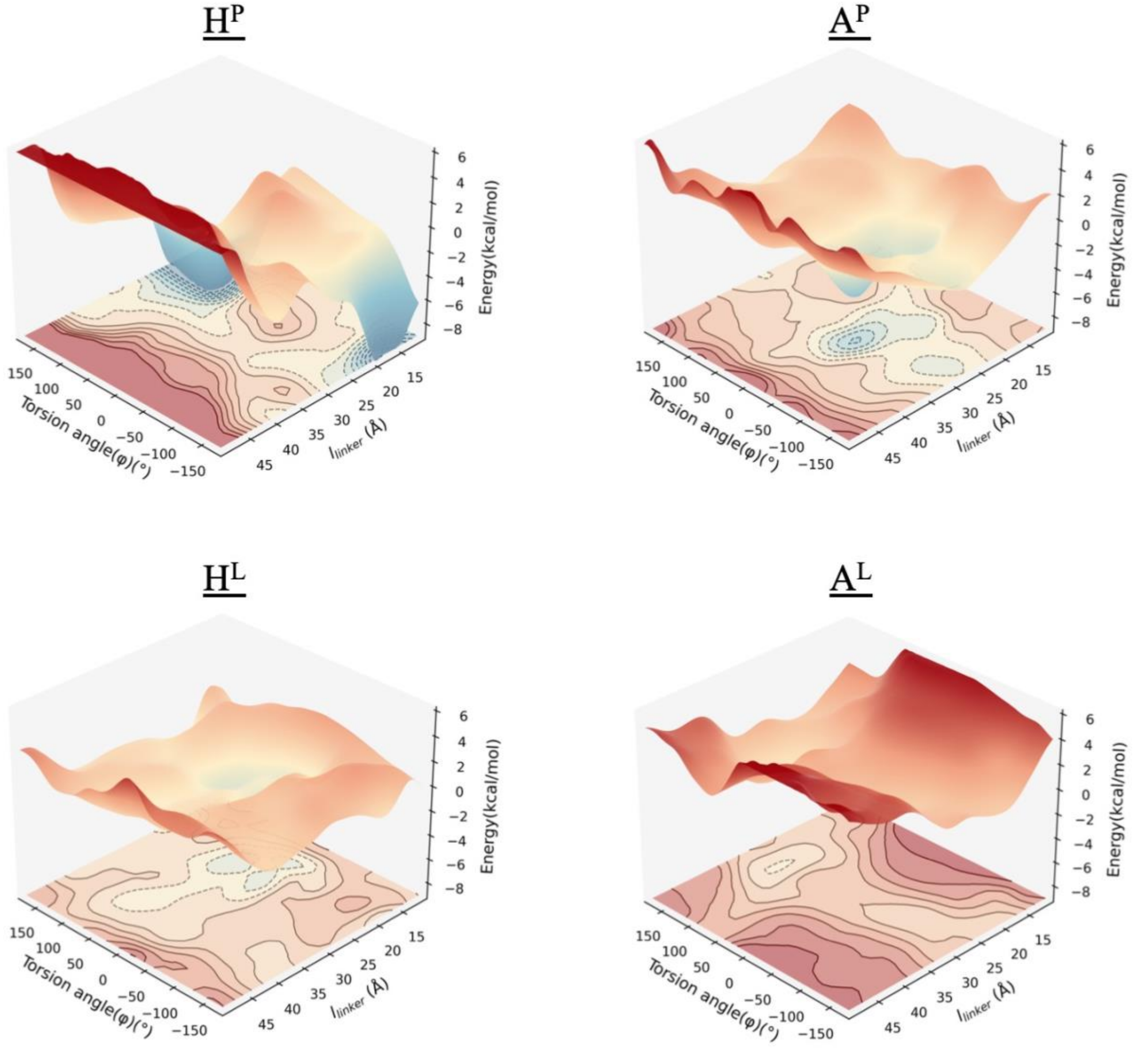


**Figure S4** | Energy landscapes sampled by MetaD simulations for *holo* CaM under **a** physiological and **b** low salt conditions, and for *apo* CaM under **c** physiological and **d** low salt conditions

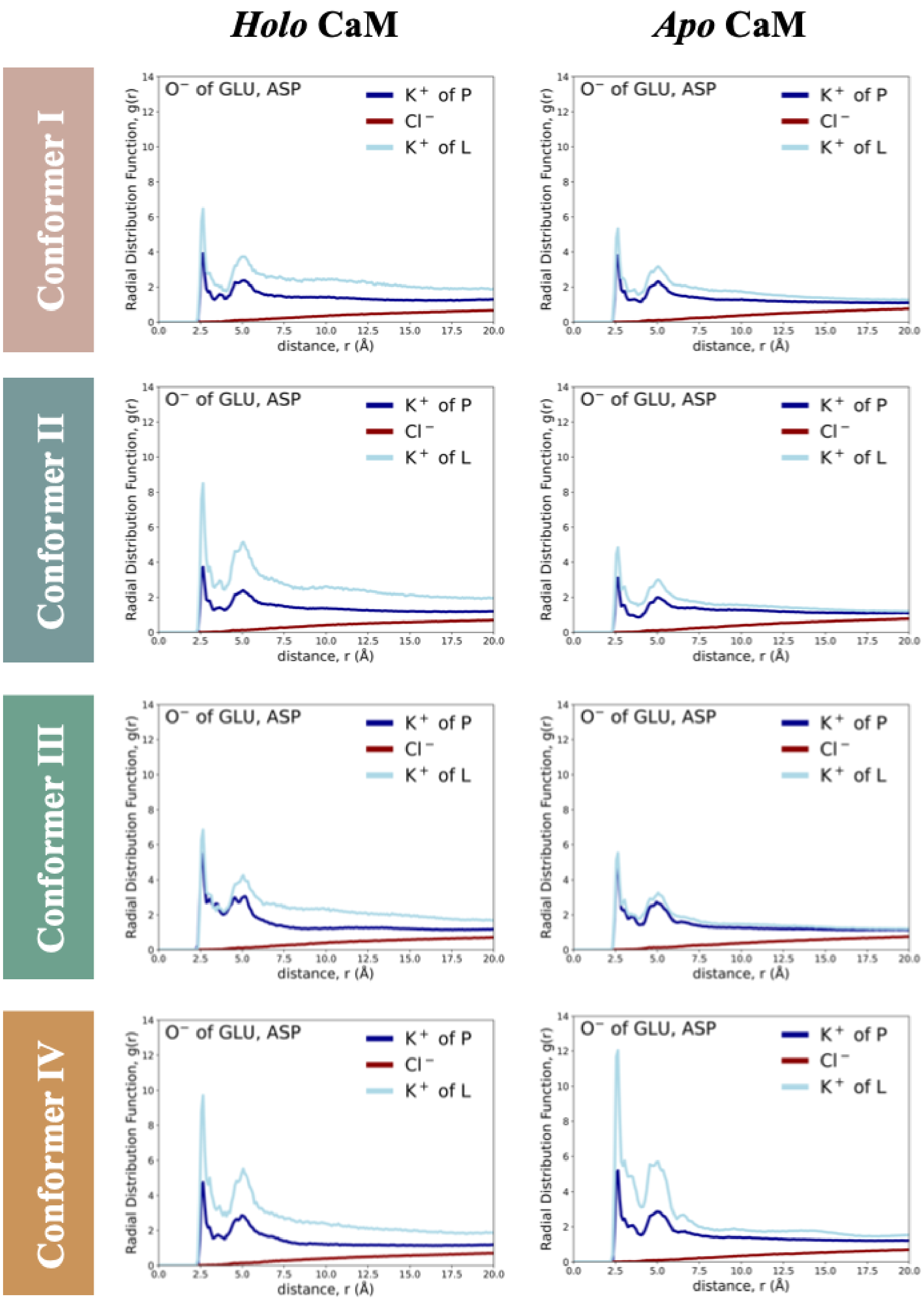


**Figure S5** | Radial distribution function, $g(r)$, between $C_\beta$ atoms of the negatively charged residues and KCl ions for all conformers. Potassium ions under low salt conditions are shown in light blue ($K^+$ of $A^L$ and $H^L$), while potassium ions under physiological salt are shown in dark blue ($K^+$ of $A^P$ and $H^P$). Chloride ions under physiological salt conditions are shown in red ($Cl^-$).

**Table S1** | Salt Bridge occupancies exceeding 20% for all conformers*

**Conformer I**

| $H^P$ Interaction | $H^P$ Occupancy (%) | $A^P$ Interaction | $A^P$ Occupancy (%) | $H^L$ Interaction | $H^L$ Occupancy (%) | $A^L$ Interaction | $A^L$ Occupancy (%) |
|---|---|---|---|---|---|---|---|
| E54-R74 | 54 | E31-K21 | 35 | E47-K77 | 58 | E31-K21 | 24 |
| E84-K115 | 36 | E47-K77 | 32 | E54-R74 | 44 | E54-R74 | 72 |
| E104-K94 | 31 | E54-R74 | 50 | E82-K77 | 50 | E82-K77 | 28 |
| E120-K75 | 23 | E87-R90 | 28 | E83-R86 | 29 | E87-R90 | 24 |
| D122-R106 | 37 | E104-K94 | 40 | E83-R90 | 25 | E104-K94 | 28 |
| | | D50-K77 | 28 | E87-R90 | 48 | E139-R86 | 20 |
| | | D78-K75 | 25 | E104-K94 | 46 | D22-K21 | 20 |
| | | D95-K94 | 24 | D95-K94 | 27 | D122-R106 | 49 |
| | | D122-R106 | 46 | D122-R106 | 61 | | |
| | | D122-R126 | 28 | D131-K30 | 39 | | |
| | | | | D133-K30 | 30 | | |

**Conformer II**

| $H^P$ Interaction | $H^P$ Occupancy (%) | $A^P$ Interaction | $A^P$ Occupancy (%) | $H^L$ Interaction | $H^L$ Occupancy (%) | $A^L$ Interaction | $A^L$ Occupancy (%) |
|---|---|---|---|---|---|---|---|
| E31-K30 | 36 | E47-K75 | 84 | E31-K21 | 26 | E31-K21 | 29 |
| E45-K30 | 22 | E54-R74 | 24 | E45-K30 | 55 | E47-K75 | 67 |
| E54-R74 | 31 | E82-K77 | 30 | E54-K75 | 77 | E54-R74 | 64 |
| E82-R86 | 22 | E82-R86 | 32 | E83-R74 | 27 | E104-K94 | 60 |
| E84-K75 | 50 | E83-R86 | 34 | E87-R90 | 22 | D22-K21 | 34 |
| E84-K77 | 24 | E104-K94 | 49 | E123-R126 | 23 | D78-K75 | 34 |
| E114-K30 | 53 | E139-R86 | 21 | E139-K75 | 33 | D122-R106 | 71 |
| D122-R106 | 22 | D122-R106 | 68 | E140-R74 | 21 | | |
| | | | | D80-R74 | 21 | | |
| | | | | D80-K75 | 27 | | |
| | | | | D122-R106 | 57 | | |

**Conformer III**

| $H^P$ Interaction | $H^P$ Occupancy (%) | $A^P$ Interaction | $A^P$ Occupancy (%) | $H^L$ Interaction | $H^L$ Occupancy (%) | $A^L$ Interaction | $A^L$ Occupancy (%) |
|---|---|---|---|---|---|---|---|
| E31-K21 | 26 | E47-K77 | 26 | E31-K21 | 45 | E31-K21 | 36 |
| E47-K77 | 47 | E54-R74 | 26 | E45-R37 | 29 | E47-K77 | 27 |
| E54-R74 | 31 | E84-K75 | 45 | E47-K77 | 45 | E54-R74 | 40 |
| E82-K77 | 20 | E87-R90 | 54 | E54-R74 | 29 | E87-R90 | 23 |
| E83-R86 | 21 | E104-K94 | 40 | E82-R86 | 20 | E104-K94 | 36 |
| E87-R90 | 33 | E139-K77 | 37 | E87-R90 | 29 | D95-K94 | 47 |
| D22-K21 | 28 | D78-K75 | 43 | E139-R74 | 22 | D122-R106 | 72 |
| D122-R106 | 78 | D80-K75 | 28 | D50-K77 | 50 | | |
| | | D95-K94 | 45 | D78-K77 | 48 | | |
| | | D122-R106 | 57 | D122-R106 | 32 | | |
| | | D122-R126 | 21 | | | | |

**Conformer IV**

| $H^P$ Interaction | $H^P$ Occupancy (%) | $A^P$ Interaction | $A^P$ Occupancy (%) | $H^L$ Interaction | $H^L$ Occupancy (%) | $A^L$ Interaction | $A^L$ Occupancy (%) |
|---|---|---|---|---|---|---|---|
| **E6-K94** | **88** | **E6-K94** | **25** | **E6-K94** | **83** | **E6-K94** | **36** |
| E7-K94 | 41 | **E47-K75** | **66** | E31-K21 | 23 | E7-K94 | 23 |
| E14-K21 | 21 | E54-R74 | 71 | **E47-K75** | **62** | E31-K21 | 48 |
| E54-R74 | 64 | E87-R90 | 49 | E54-R74 | 62 | **E47-K75** | **57** |
| **E139-K77** | **29** | E104-K94 | 26 | E87-R90 | 34 | E54-R74 | 60 |
| D78-K75 | 43 | E139-R74 | 63 | D50-R74 | 25 | E87-R90 | 22 |
| **D80-K75** | **24** | **E139-K77** | **87** | D78-K75 | 73 | E104-K94 | 41 |
| D122-R106 | 53 | D22-K21 | 23 | **D80-K75** | **43** | **E139-K77** | **81** |
| | | D78-K75 | 27 | D122-R106 | 76 | D22-K21 | 23 |
| | | D95-K94 | 28 | D122-R126 | 29 | D78-K75 | 67 |
| | | D122-R106 | 61 | | | D80-K75 | 44 |
| | | | | | | D95-K94 | 42 |
| | | | | | | D122-R106 | 47 |

* Key salt-bridge occupancies for IV that change substantially across conditions are shown in bold.
